# Genomic profiling of climate adaptation in *Aedes aegypti* along an altitudinal gradient in Nepal indicates non-gradual expansion of the disease vector

**DOI:** 10.1101/2022.04.20.488929

**Authors:** Isabelle Marie Kramer, Markus Pfenninger, Barbara Feldmeyer, Meghnath Dhimal, Ishan Gautam, Pramod Shreshta, Sunita Baral, Parbati Phuyal, Juliane Hartke, Axel Magdeburg, David A. Groneberg, Bodo Ahrens, Ruth Müller, Ann-Marie Waldvogel

## Abstract

**Background:** Driven by globalization, urbanization and climate change, the distribution range of invasive vector species has expanded to previously colder ecoregions. To reduce health-threatening impacts on humans, insect vectors are extensively studied. Population genomics can reveal the genomic basis of adaptation and help to identify emerging trends of vector expansion.

**Results:** By applying whole genome analyses and genotype-environment associations to populations of the main dengue vector *Ae. aegypti,* sampled along an altitudinal temperature gradient in Nepal (200- 1300m), we identify adaptive traits and describe the species’ genomic footprint of climate adaptation to colder ecoregions. We found two clusters of differentiation with significantly different allele frequencies in genes associated to climate adaptation between the highland population (1300m) and all other lowland populations (≤ 800 m). We revealed non-synonymous mutations in 13 of the candidate genes associated to either altitude, precipitation or cold tolerance and identified an isolation-by-environment differentiation pattern.

**Conclusion:** Other than the expected gradual differentiation along the altitudinal gradient, our results reveal a distinct genomic differentiation of the highland population. This finding either indicates a differential invasion history to Nepal or local high-altitude adaptation explaining the population’s phenotypic cold tolerance. In any case, this highland population can be assumed to carry pre-adapted alleles relevant for the species’ invasion into colder ecoregions worldwide that way expanding their climate niche.

## Background

Biodiversity, incorporating the diversity, abundance and identity of species, their genes and ecosystems, is the foundation of human health and well-being by providing essential ecosystem services. However, vector-borne diseases (VBDs), arising from inter-relationships between pathogens, invertebrate vectors and host species, also make up a part of biodiversity (1). Being detrimental to human health, one might describe VBDs and especially vector species as the dark side of biodiversity, even though vectors play also an important role in pollination (2). Annually, VBDs account for 17% of all infectious diseases worldwide (3), among those more than 390 million people are at a risk of a dengue infection (4). Worldwide, the biggest dengue virus (DENV) outbreak so far with more than 4.2 million infections was registered in 2019 (5). The current expansion of dengue fever intensified over the last decades and is predicted to further increase (6, 7). The spread of the disease via its main vector species *Aedes aegypti* (Linnaeus, 1762) was facilitated through globalisation, urbanisation and climate change (8–10). Climate warming is expected to greatly impact on the expansion processes of ectothermic insects to cooler ecoregions (9,11–13). This is not only explained by the simple fact that rising temperatures will decrease temperature barriers currently shielding cooler ecoregions thus allowing species invasion as a result of climate niche tracking (14, 15). But climate warming will furthermore rapidly move the frontier of range-edge populations thus continuously priming adaptive changes along environmental gradients (16, 17). For *Ae. aegypti* it has already been documented that populations can invade novel habitats by following their climate niches as a consequence of global warming (11, 18), moreover their expansion to new regions in the future is likely (19, 20). Further expansion to cooler ecoregions such as Europe will additionally require the adaptation to cooler temperatures (21, 22). It is, however, less clear whether range-edge populations carry sufficient adaptive potential for further acceleration of their expansion process.

Climatic clines influence population divergence as shown in *Anopheles gambiae* (23), *Drosophila melanogaster* (24–26) and recently for the first time in *Ae. aegypti* (27). Invasive species that experience range expansion along such clines are expected to locally adapt (28). Genetic admixture can benefit invaders by either mitigating the negative effects of bottlenecks during their introduction by masking deleterious alleles and/or by generating new allelic combinations causing many phenotypes, which provides raw material for selection and rapid adaptation (29). For instance, *Ae. albopictus* adapted genetically and morphometrically to Northern latitudes prior to its successful worldwide expansion (22) and *Drosophila melanogaster* preadapted to the temperate and tropical conditions that they then encountered in North America and/or Australia prior to their invasion (24). Thus, genomic signatures of ‘climate adaptation’ are a special case of classical local adaptation, since environmental heterogeneity or ideally the gradual variation of climate along environmental gradients will result in gradual or at least environmentally correlated signatures of selection (30). Climate and local adaptation of *Ae. aegypti* to colder climates is scarcely investigated, consequently Schmidt and colleagues (31) recently identified the need to better investigate adaptive traits and associated gene sets in mosquito species. Population genomics is thus a straightforward approach to examine the influence of climate on adaptation in various organisms (15).

To recognize emerging trends in adaptive traits of *Ae. aegypti* to cooler ecoregions driven by climate warming, the study of *Ae. aegypti* currently spreading along climatic transects with ongoing disease expansion (e.g. Dengue) in the Hindu Kush Himalayan (HKH) country Nepal could provide useful insights (32–41). After an introduction into a new environment, populations are unlikely at their fitness optimum at first and, therefore, adapt to new conditions through environmental selection (27). According to this theory, the initial overwintering potential in a highland population (1300m) can be lower compared to the lowlands (≤ 800 m; 21,42). However, Nepal is suffering under climate warming that influences the climate along altitudinal gradients extremely (41,43; unpublished data- Phuyal, Kramer et al. 2021), not only with regard to temperature but likely also humidity (44). Maldaptation of the mosquitos to higher altitudes might thus be facilitated by climate change. We thus tested if gradual climate heterogeneity along an altitudinal gradient in Central Nepal is reflected in patterns of genomic differentiation of natural *Ae. aegypti* populations sampled along the gradient, henceforth referred to as pattern of ‘climate adaptation’. Here, climate adaptation is studied in *Ae. aegypti* field populations sampled along a prominent climate gradient of Nepal in the mountain region, using the currently most commonly applied genotype-environment association (GEA) tool (LFMM, Figure 1; 45).

**Figure 1.**
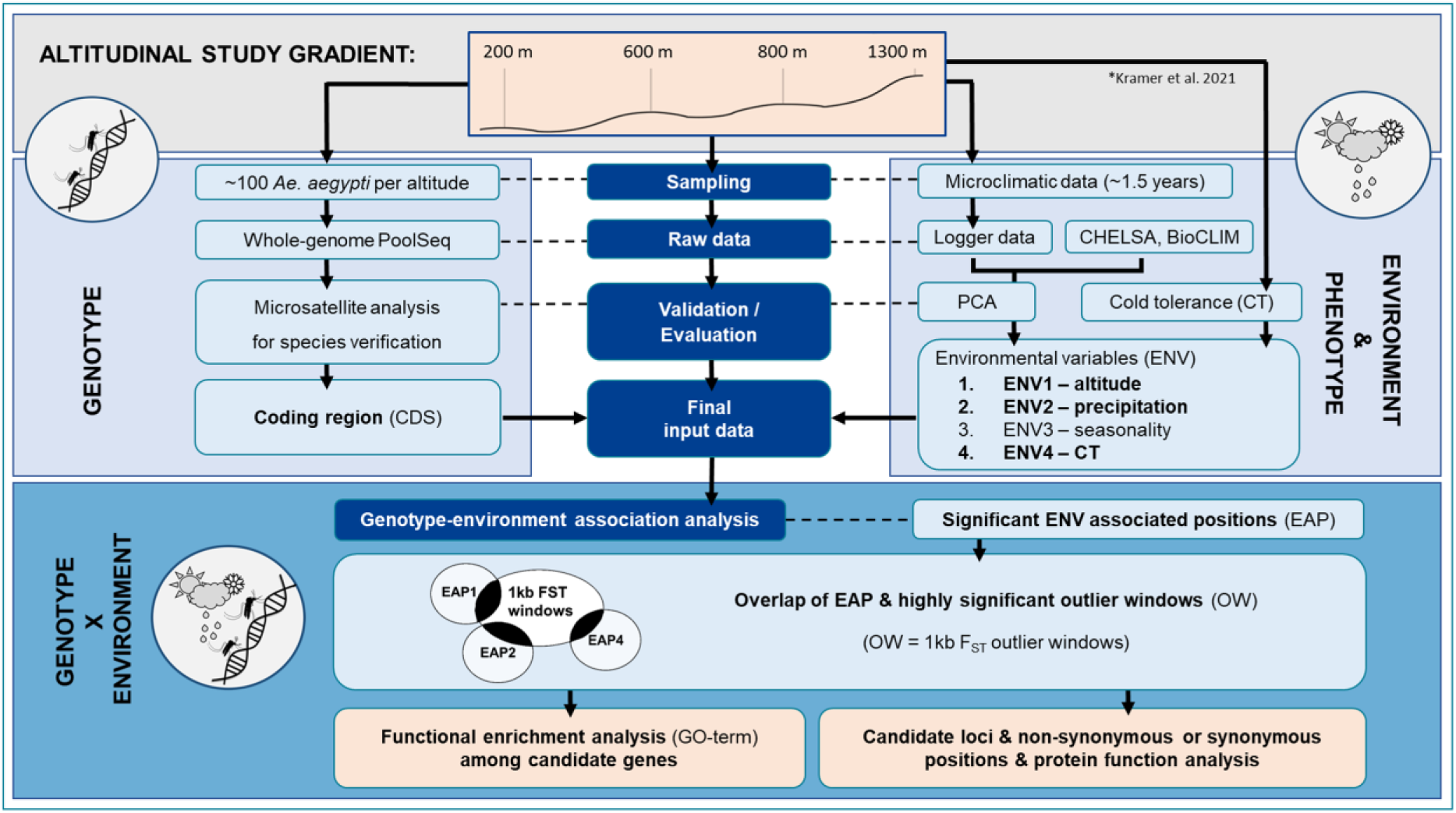
Study design to analyze climate adaptation of natural *Ae. aegypti* populations along an altitudinal gradient. CDS= Coding region, CT= cold tolerance data normalized to controls; (42), ENV= environmental variable, EAP= significant ENV associated positions, OW= significant 1kb F_ST_ outlier windows.

## Results

In total, we collected 1) four *Ae. aegypti* populations from Chitwan (CH200, 200 m above sea level), Dhading (DH600, 600 m asl), Dharke (DK800, 800 m asl) and Kathmandu (KT1300, 1300 m asl; (42, 46), and 2) high-resolution microclimate data, open access weather data (CHELSA) and phenotypic expression of study populations (experimental cold tolerance data, Figure 1; (42). We first confirmed that all Nepalese populations belong to one *Ae. aegypti* subspecies using a µsats analysis and second by means of a population genomic approach (Pool-Seq) and subsequent GEA analysis (LFMM), we identify 33 candidate genes for climate adaptation containing non-synonymous or synonymous mutations, and discuss their functional basis by conducting a literature survey. In addition, 1200 candidate genes for local adaptation were identified, among them known loci involved in insecticide resistance (knockdown resistance (*kdr*) mutations (V1016G, F1534C, and S989P)) and metabolic resistance (47) and vector competence.

### Subspecies analysis

The STRUCTURE analysis based on µsats extracted from our Nepalese Pool-Seq data confirms that our genomic data sets only consist of one *Ae. aegypti* subspecies. All of the ten runs with STRUCTURE using K = 2 with six µsats in a comparison to other populations worldwide (West Africa, Costa Rica, Australia- Innisfail; (48)) indicate that the African population is different from the Nepalese populations, the Costa Rica population is similar to the Nepalese populations and the Australian is similar to the African population (Figure 2a, Additional file 1 Figure 1). Due to lower coverage of the population from KT1300 of Nepal, only five µsats were included (AC1 excluded) and the Australian population was restricted to three µsats (A9, AC1 and B3 were excluded). When comparing Nepalese populations amongst each other (10/10 runs with K2-11 µsats), low similarities between the CH200 and KT1300 population and higher similarities between the CH200, DH600 and DK800 populations are present (Figure 2b, Additional file 1 Figure 1). K = 3 displays that the lowest sampling sites (CH200 and DH600) show similarities in comparison to the populations from higher altitude and K = 4 shows a distinct structural difference between the four populations (Figure 2b, Additional file 1 Figure 1).

**Figure 2.**
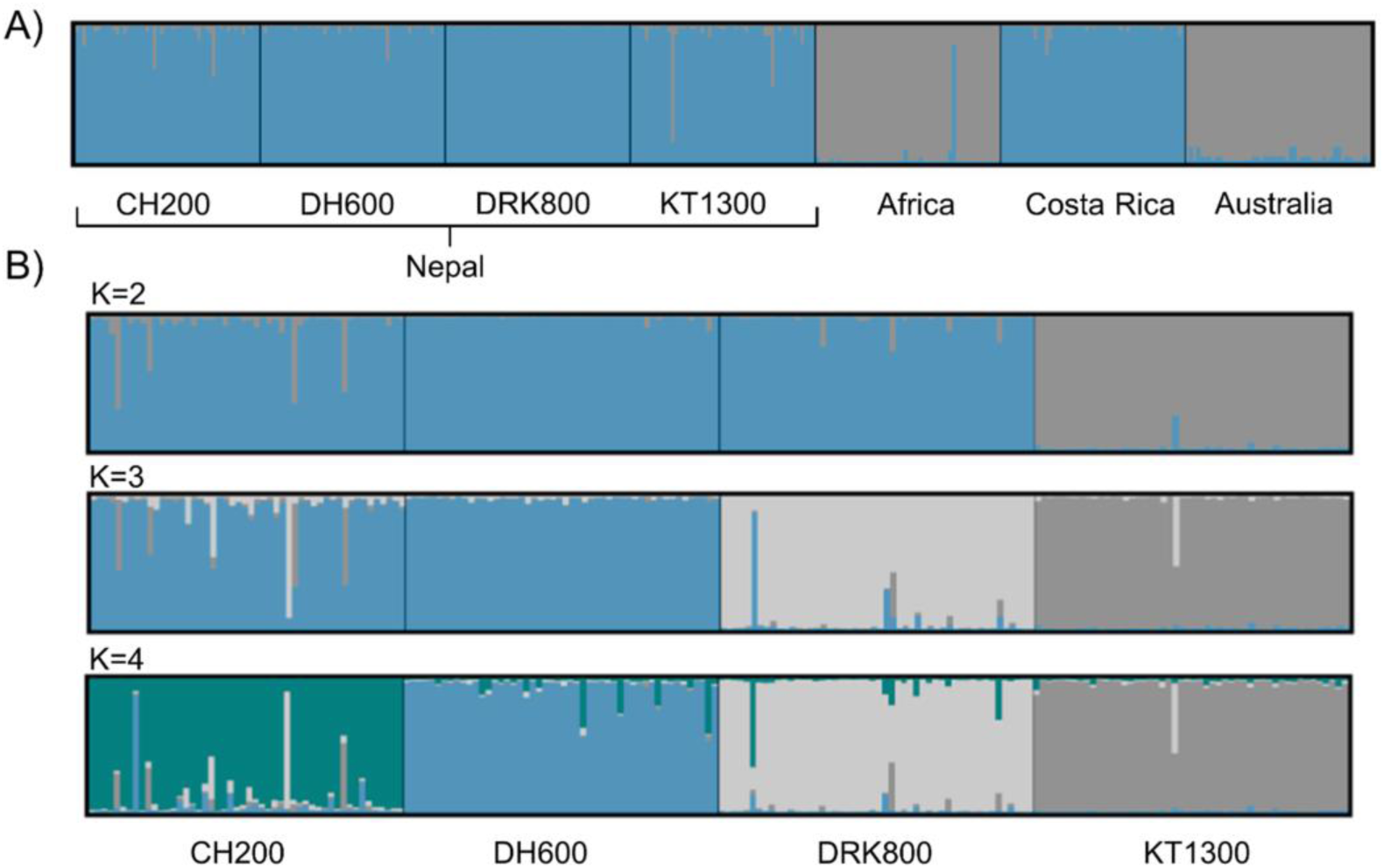
Global (A) and local (B) genetic structure of *Aedes aegypti* populations. Comparison of four populations from Nepal A) with populations from Africa, Costa Rica and Australia (48) using 6 microsatellite regions (K=2) and B) with each other using 11 microsatellite regions (K=2-4; Additional file 1 Figure 1). Altitude of sampling sites of *Ae. aegypti* populations in Central Nepal: CH200 = 200 m asl (Chitwan), DH600 = 600 m asl (Dhading), DK800 = 800 m asl (Dharke), KT1300 = 1300 m asl (Kathmandu).

### Population differentiation

Nucleotide diversity (π) is smaller in exonic regions compared to the genome-wide average (per site) and all populations show a similarly low π with an average of 0.0127 in 1kb windows. The low-altitude population CH200 has the highest population mutation rate (θ), however, there is no increasing trend towards higher altitude. Concerning the effective population size (Ne), there is a trend towards decreasing values along the altitudinal gradient, however smallest Ne is found in DH600 (Table 1).

**Table 1.** Mapping and coverage statistics of four *Ae. aegypti* populations sampled along an altitudinal gradient. Population genomic parameters estimated per site (1b) or in non-overlapping 1kb-windows: nucleotide diversity (π), population mutation parameter theta (θ) and effective population size (Ne) calculated as Ne= θ/4µ with µ= 2.1 × 10−9 (108). Altitude of sampling sites of *Ae. aegypti* populations in Central Nepal: CH200 = 200 m asl (Chitwan), DH600 = 600 m asl (Dhading), DK800 = 800 m asl (Dharke), KT1300 = 1300 m asl (Kathmandu).

| Parameter | region | window | CH200 | DH600 | DK800 | KT1300 |
| --- | --- | --- | --- | --- | --- | --- |
| mapped reads (%) | genome- wide | - | 66.07 | 66.79 | 64.27 | 65.99 |
| mean coverage | genome- wide | - | 22.46 | 17.86 | 21.1 | 19.02 |
| genome coverage (%) | genome- wide | - | 72.66 | 63.87 | 70.64 | 66.55 |
| PoPoolation analysis |  |  |  |  |  |  |
| $\pi$ | genome- wide | 1kb | 0.0130 | 0.0125 | 0.0129 | 0.0126 |
|  | genome- wide | 1b | 0.0135 | 0.0132 | 0.0135 | 0.0133 |
|  | exon | 1b | 0.0079 | 0.0077 | 0.0080 | 0.0077 |
| $\theta$ | genome- wide | 1kb | 0.0130 | 0.0125 | 0.0129 | 0.0127 |
| $N_e$ | genome-wide | 1kb | 1,548,452.4 | 1,485,238.1 | 1,536,904.8 | 1,507,976.2 |

Mean pairwise FST range between 0.05-0.067 (Table 2) indicating low levels of genomic differentiation and high relatedness among the four Nepalese populations (Figure 3, Additional file 1 Figure 2), in line with the results from the µsats analysis (Figure 2). Moreover, the Mantel test revealed no signs of isolation by distance (p=0.67, r=-0.27).

**Figure 3.**
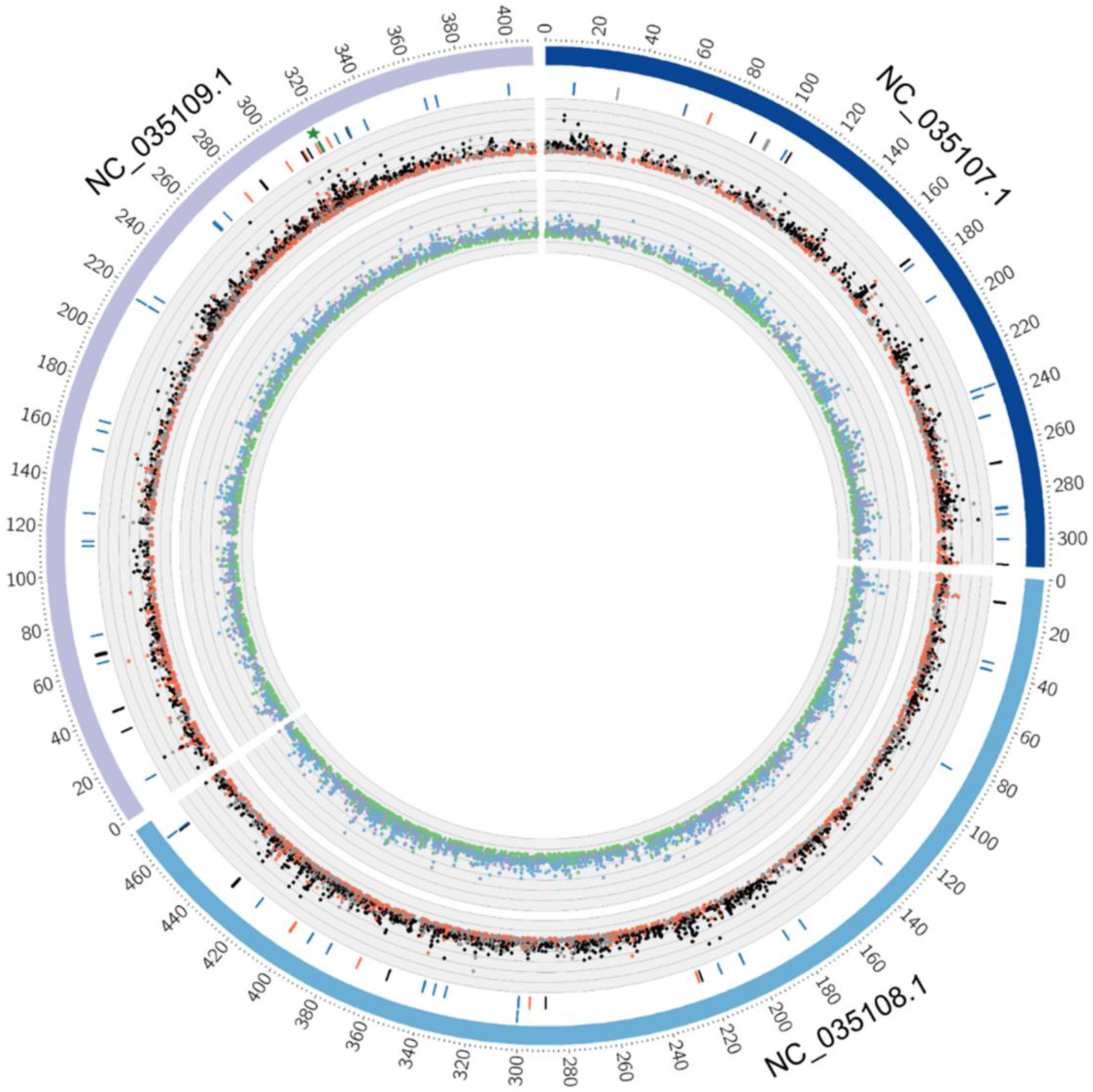
Genome wide pairwise F_ST_ distribution per 1kb-windows (OW) of Nepalese *Ae. aegypti* populations. The three chromosomes of the *Ae. aegypti* genome are represented in the outermost circle. From innermost to outermost circle: (A) the innermost circle shows the pairwise F_ST_ distribution (range:0-0.7) in 1kb windows between the lowland populations (purple: CH200 vs. DH600; green: CH200 vs. DK800; light-blue: DH600 vs. DK800), (B) the middle circle shows the comparison between the lowland populations and the KT1300 (red: CH200 vs. KT1300; black: DH600 vs. KT1300; grey: DK800 vs. KT1300), (C) The white circle gives the position of all EAP-OW genes (black), the candidate genes containing non- synonymous mutations (red), the detoxification genes containing significant positions (blue), the voltage-gated sodium channel (green and a green star) and the vector competence genes (grey). Altitude of sampling sites of *Ae. aegypti* populations: CH200 = 200 m asl, DH600 = 600 m asl, DK800 = 800 m asl, KT1300 = 1300 m asl.

**Table 2.** Mean F_ST_ value and number of candidate SNPs/genes for climate and local adaptation. Altitude of sampling sites of *Ae. aegypti* populations: CH200 = 200 m asl, DH600 = 600 m asl, DK800 = 800 m asl, KT1300 = 1300 m asl. Climate adaption: Outline of stringent signatures of climate selection (ENV1 ∼ altitude, ENV2 ∼ precipitation, ENV4 = cold tolerance) by overlapping outlier windows of highly significant population differentiation (OW; 1kb- window) with EAPs (GEA gene list) for each population comparison. Local adaptation: overlap between F_ST_ 1kb-window outlier analysis (OW) and the F_ST_ 1b-window outlier analysis (OP) excluding SNPs and genes from the EAP-OW analysis (OW-OP). Numbers given per position and per gene hit: integrated-hits/unique-hits/non-synonymous. If not marked otherwise, all unique-hits are also present in the upper 1% tail of the site-specific F_ST_ distribution (OP).

|  | CH200-DH600 | CH200-DK800 | CH200-KT1300 | DH600-DK800 | DH200-KT1300 | DK800-KT1300 | Total |
| --- | --- | --- | --- | --- | --- | --- | --- |
| mean $F_{ST}$ | 0.05784 | 0.04996 | 0.05567 | 0.06251 | 0.06676 | 0.05822 | - |
| **OW-OP** |  |  |  |  |  |  |  |
| Highly significant site-specific positions | 1251 | 1171 | 1250 | 1331 | 1400 | 1189 | 6303 |
| Number of genes (without duplicate) | 277 | 280 | 284 | 269 | 286 | 263 | 1200 |
| **OW** |  |  |  |  |  |  |  |
| Highly significant 1kb-windows | 2919 | 2911 | 2928 | 2940 | 2931 | 2922 | - |
| Number of genes (without duplicates) | 535 | 576 | 579 | 501 | 551 | 550 | - |
| **Overlap of OW and EAP- number of positions** |  |  |  |  |  |  |  |
| ENV1 | 0 | 0 | 4 | 1 | 4 | 2 | 11/9/2 |
| ENV2 | 2 | 2 | 19 | 1 | 15 | 20 | 59/40*/12 |
| ENV4 | 0 | 0 | 8 | 1 | 7 | 4 | 20/14/3 |
| **Overlap of OW and EAP- number of genes** |  |  |  |  |  |  |  |
| ENV1 | 0 | 0 | 5 | 1 | 3 | 1 | 10/8 |
| ENV2 | 2 | 2 | 15 | 1 | 13 | 14 | 47/31 |
| ENV4 | 0 | 0 | 8 | 1 | 6 | 3 | 18/13 |

\*9 positions are not present in the upper 1% tail of the site-specific $F_{ST}$ distribution, including five non-synonymous SNPs

### Environmental data

The annual and seasonal CHELSA data shows a gradual decrease of mean, minimum and maximum temperature along the altitudinal gradient (Additional file 1 Figure 3). The microclimate data shows higher variability throughout the seasons with a decreasing trend of mean and minimum temperature with increasing altitude but higher variability in the maximum temperature, especially at DH600 (Additional file 1 Figure 4). CHELSA data reveals a precipitation pattern similar for all sampling sites (Additional file 1 Figure 5). To reduce confounding covariation in the environmental data set a principal component analysis (PCA) was run. The first three components of the PCA are mainly related to the following environmental factors: PCA1 – altitude (70.86%; ENV1), PCA2 – precipitation (27.45%; ENV2) and PCA3 – seasonality (1.69%; ENV3; Additional file 1 Table 5, Additional file 1 Figure 6-9).

**Figure 4.**
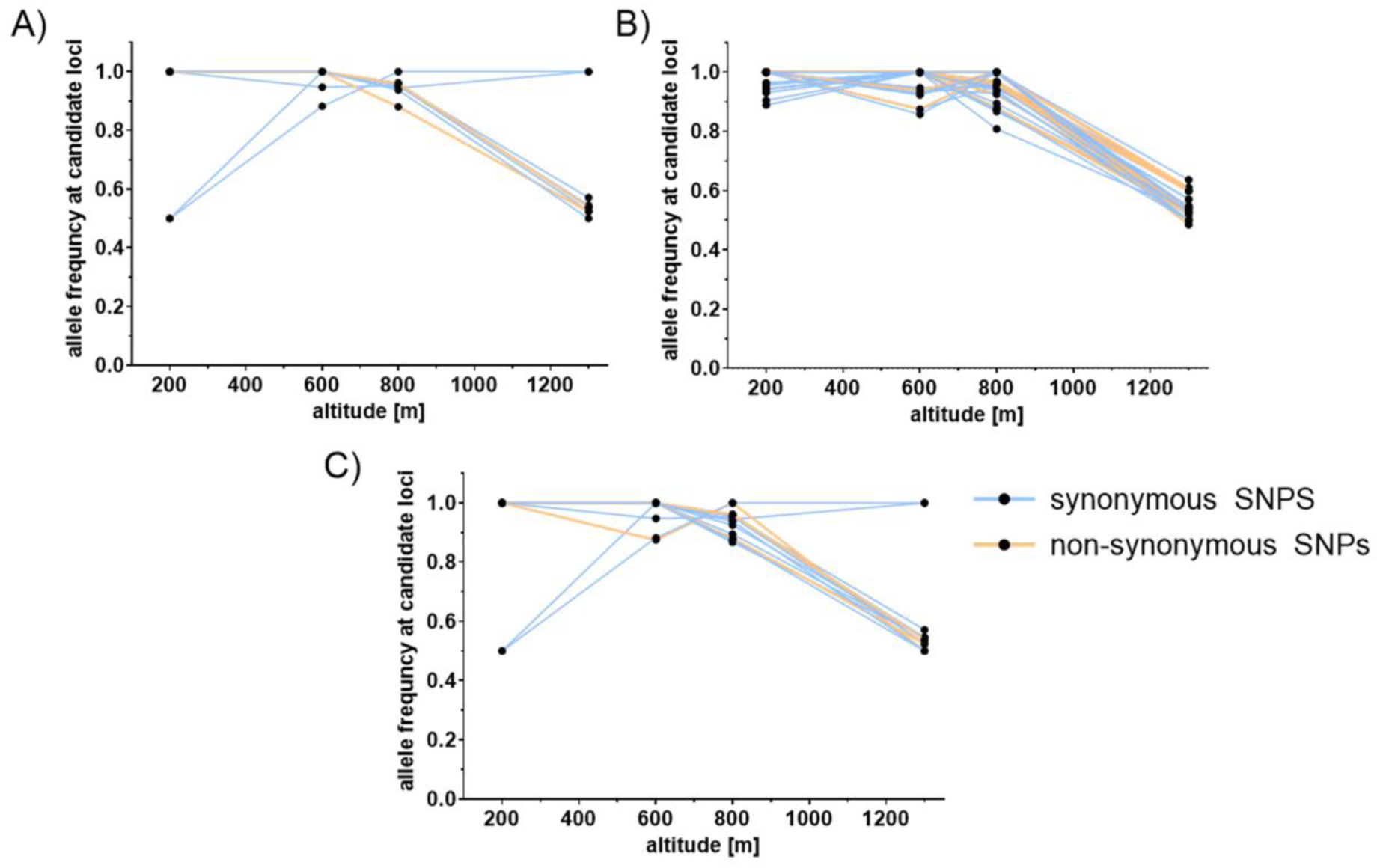
Allele frequencies of candidate loci (EAP-OW) plotted against the altitudinal gradient of their population origin. Candidate loci associated with A) ENV1 ∼ altitude, B) ENV2 ∼ precipitation, C) ENV4 = cold tolerance. Details on non-synoynomous SNPs in Table 3.

**Figure 5.**
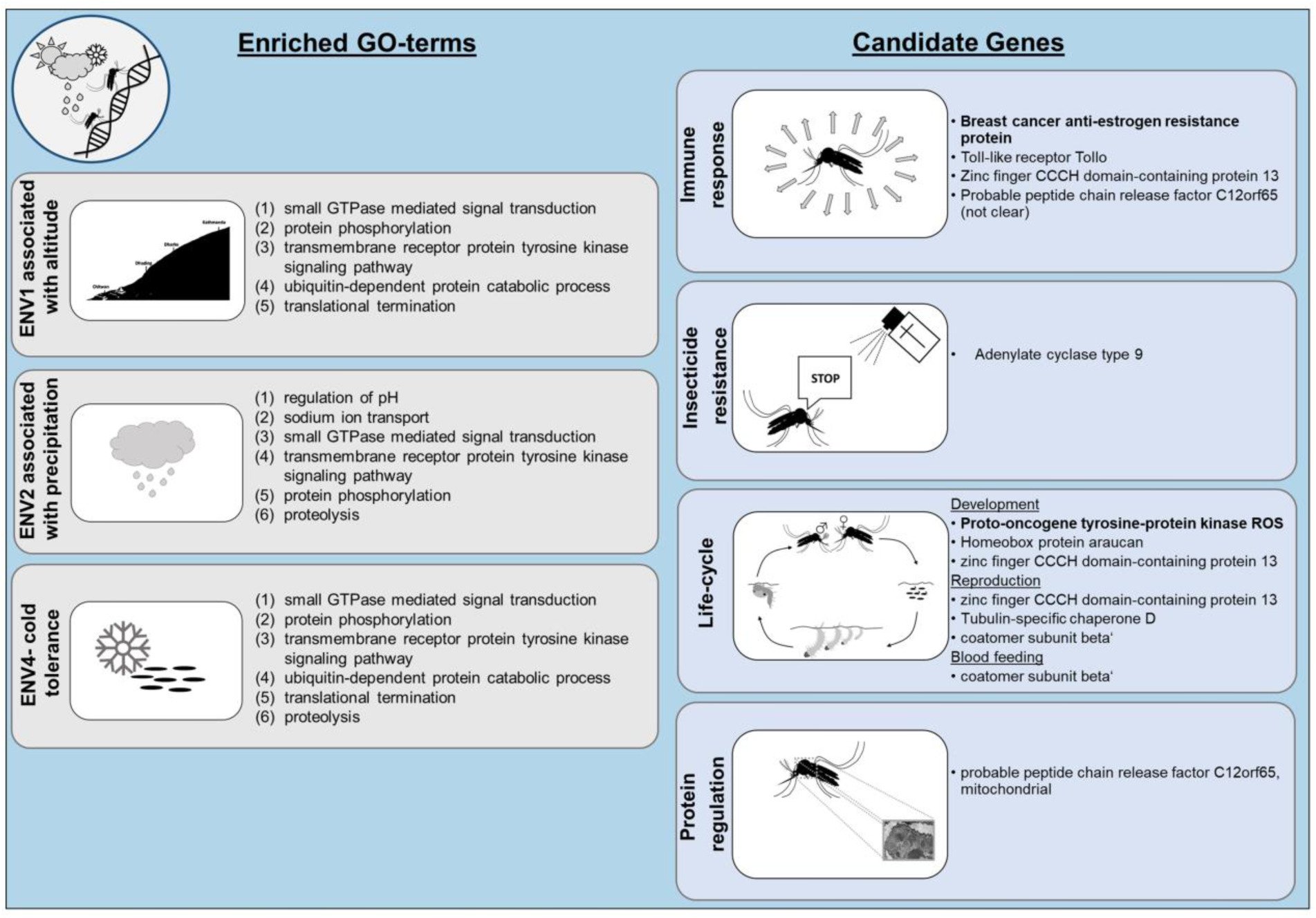
Significantly enriched GO terms for each environmental variable and candidate genes sorted by functional groups (see Table 4). Candidate genes only carrying non-synonymous mutations are given. Genes written in bold are associated with all three environmental variables (ENV), whereas all the others are only associated with precipitation. Three uncharacterized genes given in Table 3 are not shown.

**Figure 6.**
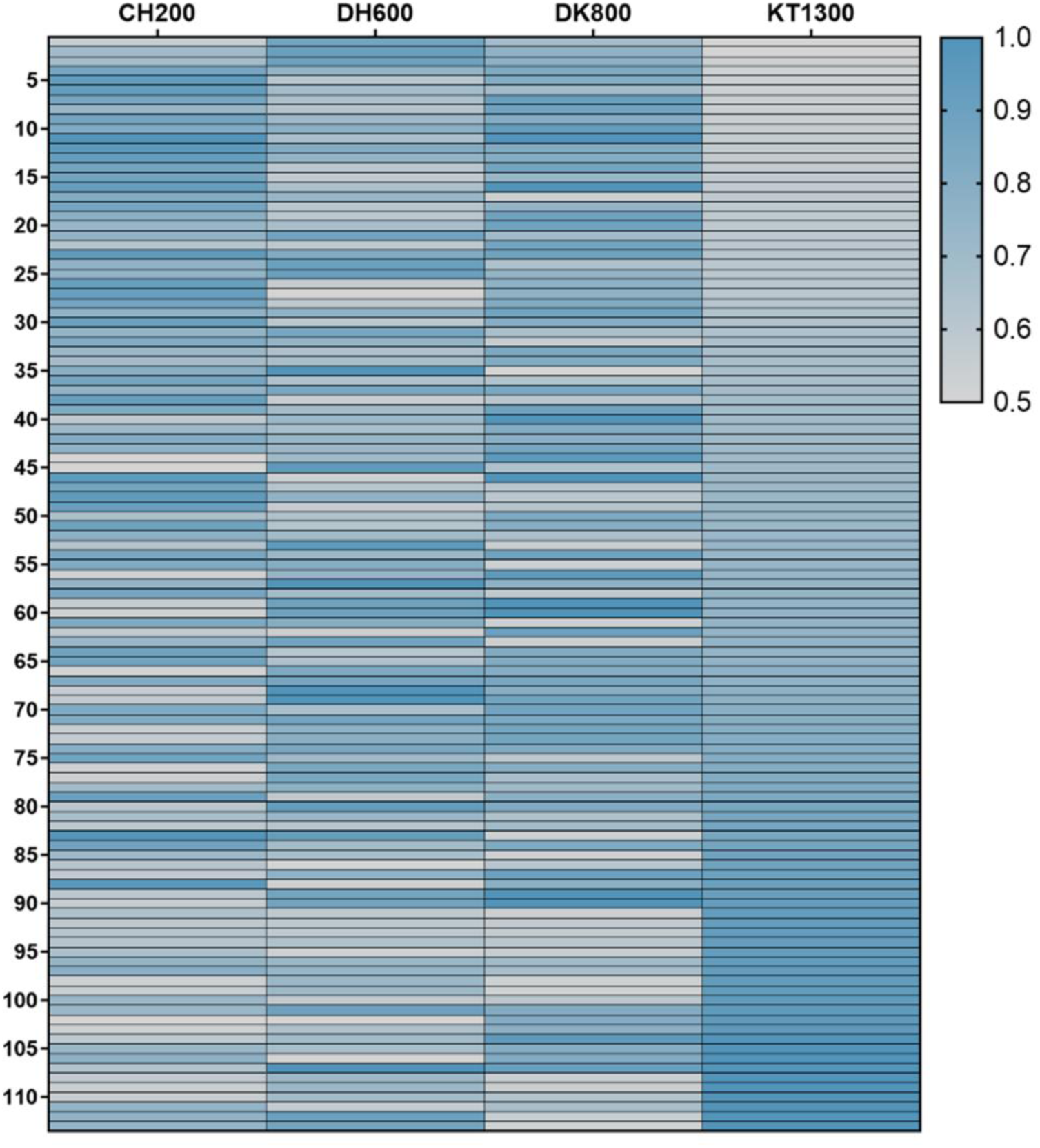
Heat map of allele frequency distribution at candidate loci containing non- synonymous mutations. In total 113 detoxification genes of *Ae. Aegypti* are given. Allele frequencies were sorted after KT1300. Altitude of sampling sites of *Ae. aegypti* populations: CH200 = 200 m asl, DH600 = 600 m asl, DK800 = 800 m asl, KT1300 = 1300 m asl.

**Table 3.**
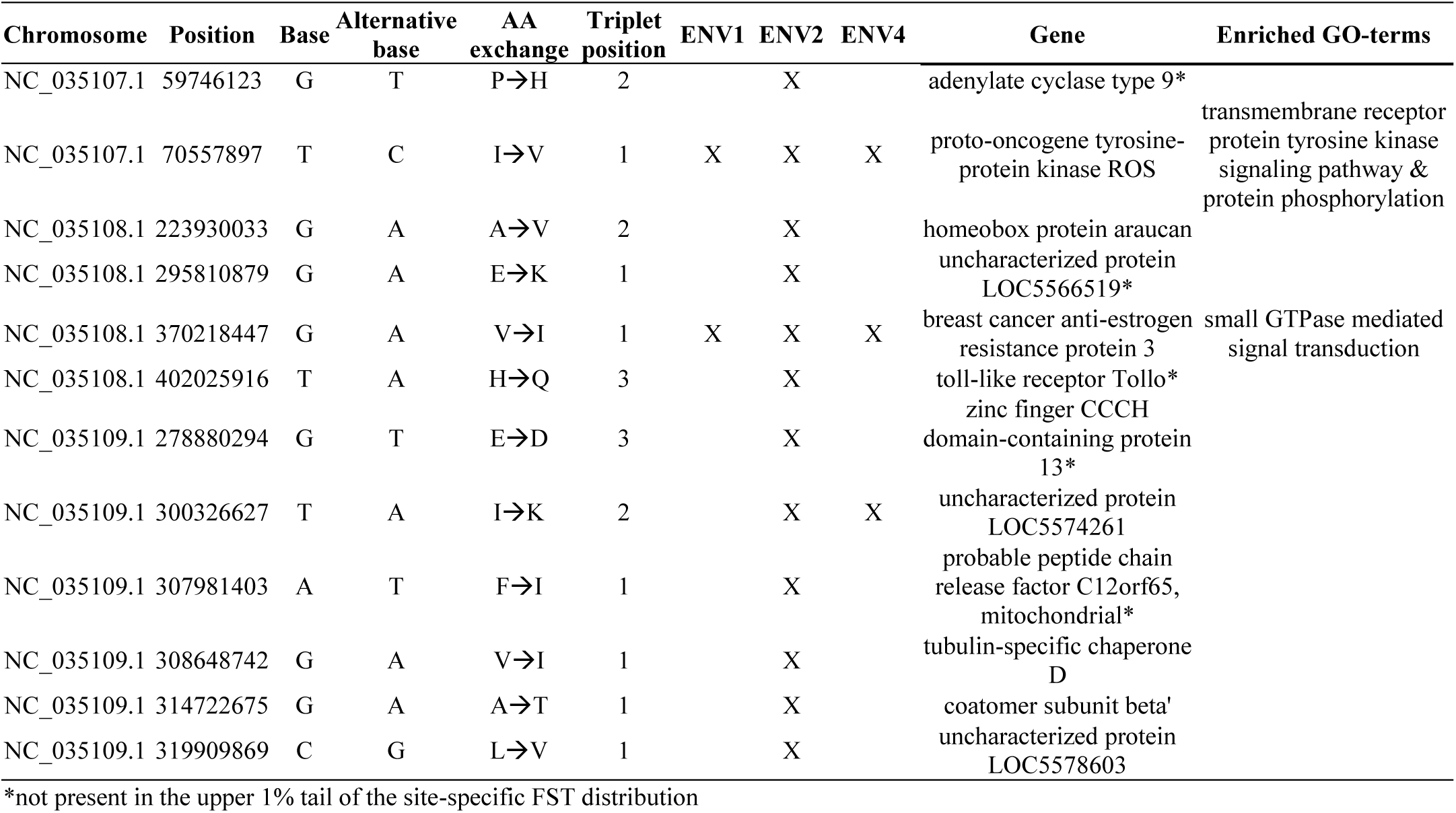
Non-synonymous substitutions of EAP-OWs that indicate significant involvement of genes in climate adaptation. The genomic position, base, alternative base, amino acid (AA) exchange, association to respective environmental variables (ENV1 ∼ altitude, ENV2 ∼ precipitation, ENV4 = cold tolerance) and the annotated candidate gene are given. Significantly enriched GO-terms (Figure 5) are mentioned if they can be linked to the candidate gene using uniprot.

**Table 4.**
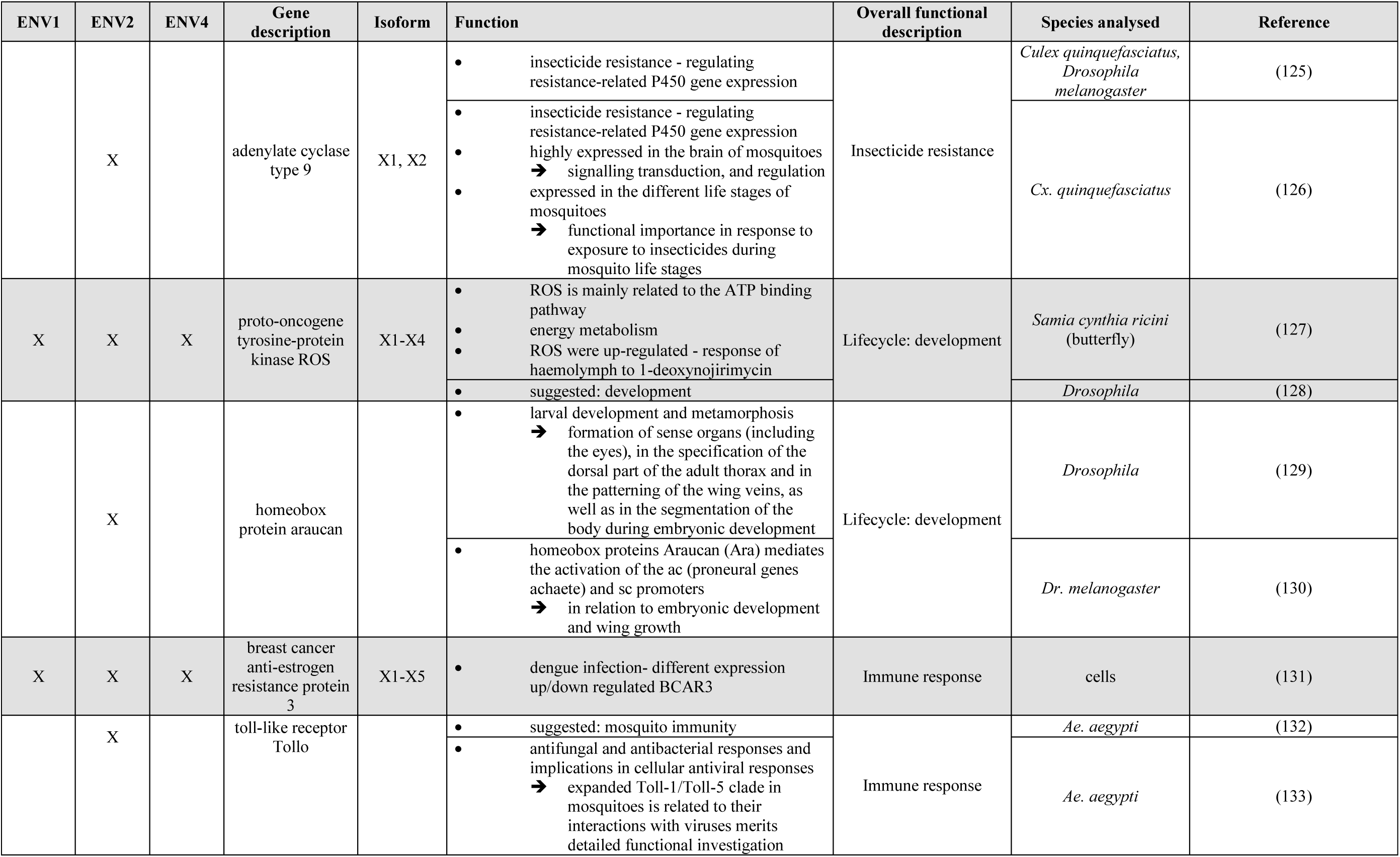
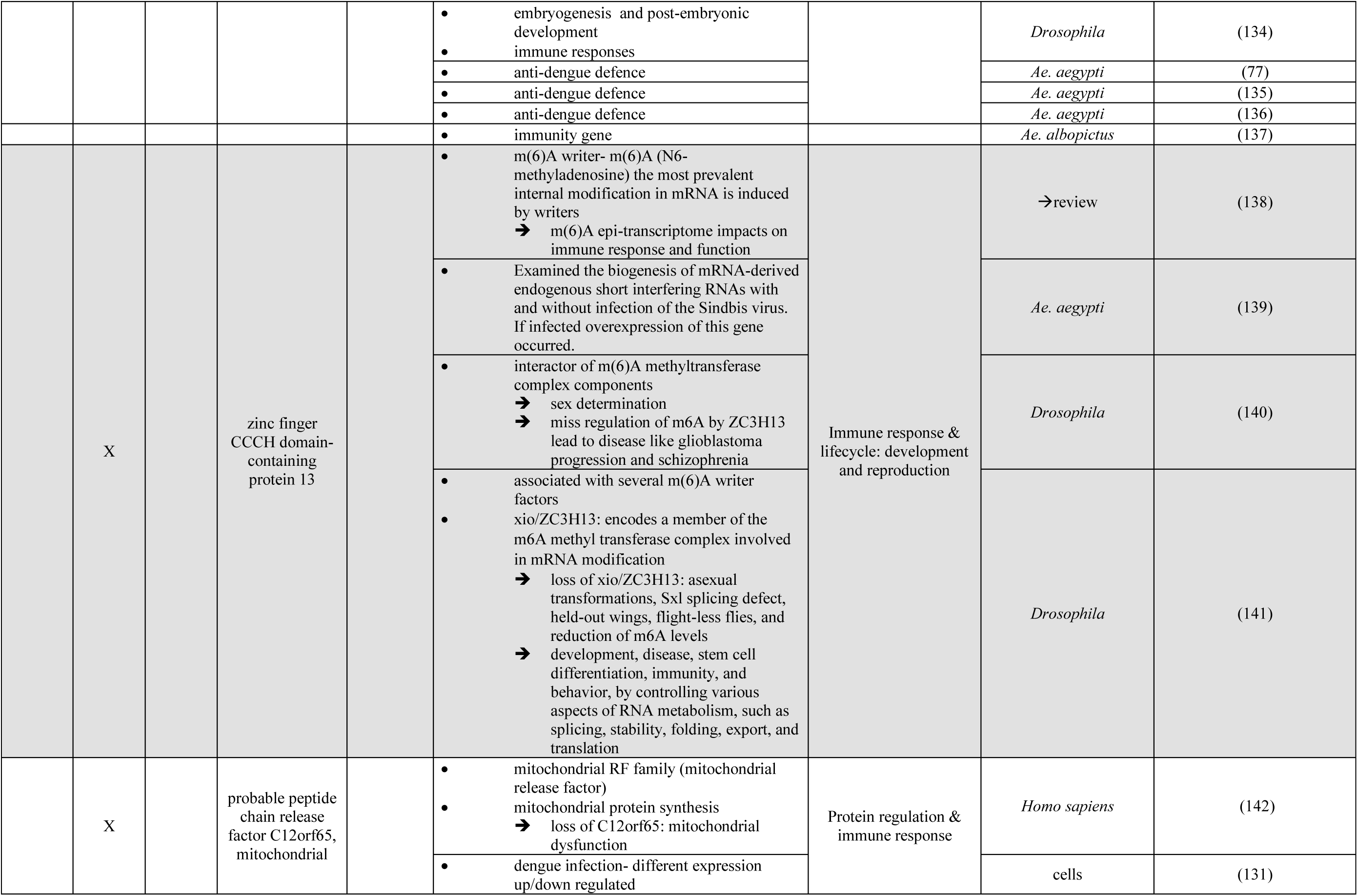
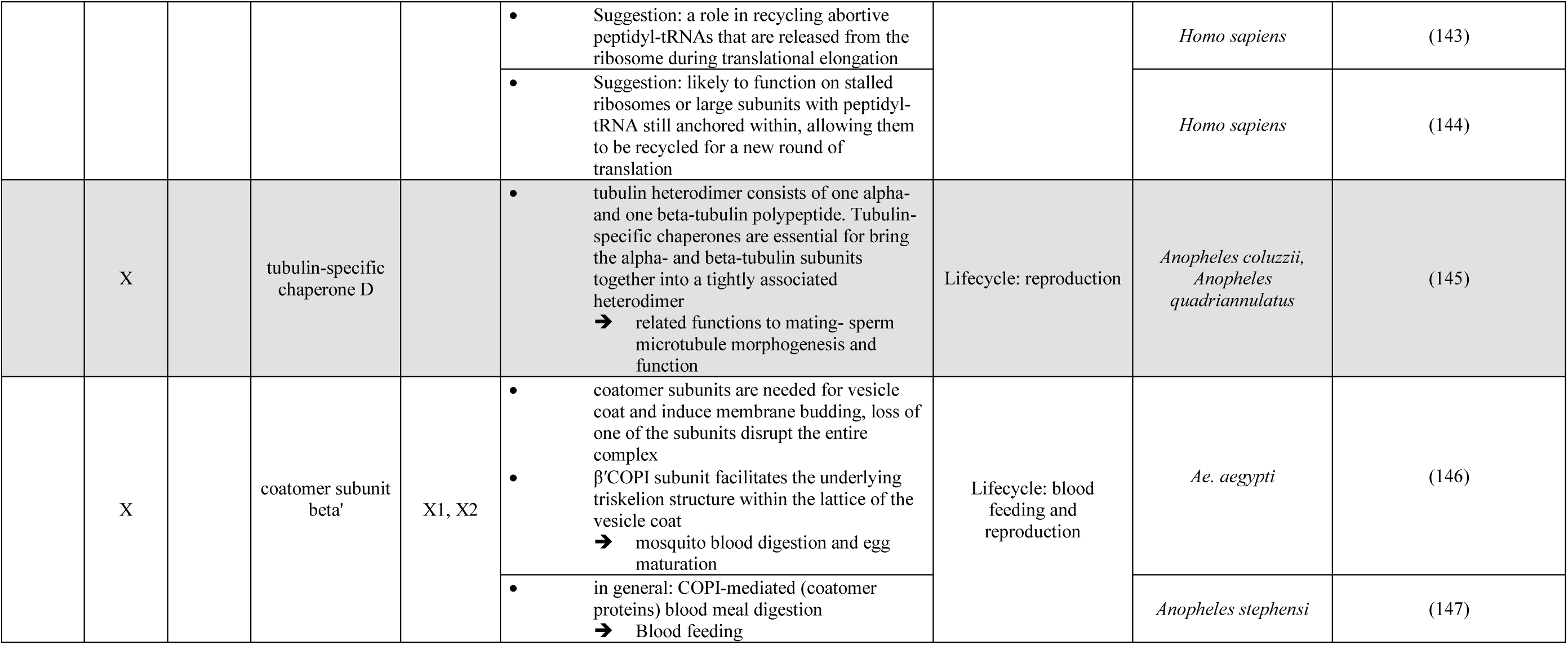
Details on gene function of the nine characterized candidate genes associated to environmental variables. ENV1 ∼ altitude. ENV2 ∼ precipitation, ENV4 = cold tolerance. Three other uncharacterized genes are not included in this list.

**Table 5.** Allele frequency and allelic variant of *kdr* mutations with exact genome position. Altitude of sampling sites of *Ae. aegypti* populations: CH200 = 200 m asl, DH600 = 600 m asl, DK800 = 800 m asl, KT1300 = 1300 m asl.

| Allele frequency |  |  |  |  |  |  |  |
| --- | --- | --- | --- | --- | --- | --- | --- |
| Mutation ID | Chromosome | Position | Amino acid code | CH200 | DH600 | DK800 | KT1300 |
| S989P | NC_035109.1 | 315984077 | TCC – wildtype | 1 | 1 | 1 | 1 |
|  |  |  | CCC – mutant | 0 | 0 | 0 | 0 |
| V1016G | NC_035109.1 | 315983762 | GTA – wildtype | 0.68 | 0.93 | 0.78 | 0.56 |
|  |  |  | GGA – mutant | 0.32 | 0.07 | 0.22 | 0.44 |
| F1534C | NC_035109.1 | 315939224 | TTC – wildtype | 0.94 | 0.88 | 0.91 | 0.69 |
|  |  |  | TGC – mutant | 0.06 | 0.12 | 0.09 | 0.31 |
| Allelic variant (total count) |  |  |  |  |  |  |  |
| S989P | NC_035109.1 | 315984077 | TCC – wildtype | 26 | 22 | 23 | 26 |
|  |  |  | CCC – mutant | 0 | 0 | 0 | 0 |
| V1016G | NC_035109.1 | 315983762 | GTA – wildtype | 15 | 14 | 14 | 10 |
|  |  |  | GGA – mutant | 7 | 1 | 4 | 8 |
| F1534C | NC_035109.1 | 315939224 | TTC – wildtype | 17 | 15 | 10 | 9 |
|  |  |  | TGC – mutant | 1 | 2 | 1 | 4 |

### Genotype-environment association

The LFMM analysis reveal 47 single nucleotide polymorphisms (SNPs) within 46 genes associated to ENV1 (associated with altitude), 216 SNPs within 172 genes associated to ENV2 (associated with precipitation), zero SNPs associated to ENV3 (associated with seasonality) and 69 SNPs within 64 genes associated to ENV4 (cold tolerance) (Table 2; Additional file 1 Figure 10). After our stringent filtering when overlapping significant ENV associated positions (EAPs) with highly significant FST outlier windows (OW; 1 kb-window; Figure 1) 9 SNPs within 8 genes associated to altitude (ENV1) are present. We accordingly retain 40 SNPs within 31 genes associated to precipitation (ENV2) and 14 SNPs within 13 genes associated to cold tolerance (ENV4; Table 2, Additional file 1 Figure 2, Figure 3). All EAP- OW (overlap of EAP with OW) SNPs are also present in highly significant outlier positions per site (OP) except 9 SNPs associated with precipitation (Table 2; Additional file 2 Table 2- Table 4). Observed allele frequencies plotted against the altitudinal gradient of population origins do not support the expected gradual variation of allele frequencies at candidate positions (EAP-OW). Instead of a pattern of gradual variation, our results reveal a major difference in allele frequency of candidate loci in KT1300 compared to all other lowland populations (CH200, DH600, DK800; Figure 4).

### Functional enrichment associated to climate adaptation

The investigated populations of *Ae. aegypti* across the Nepalese altitudinal gradient reveal 33 candidate genes with signatures of climate selection (temperature (ENV1): 8, precipitation (ENV2): 31, cold tolerance (ENV4): 13), which equals to ∼0.2% of protein-coding genes with signatures of climate selection. Functional analysis of the eight genes that are associated with altitude (ENV1) yielded the following five GO terms to be significantly enriched: 1) ‘small GTPase mediated signal transduction’, 2) ‘protein phosphorylation’, 3) ‘transmembrane receptor protein tyrosine kinase signaling pathway’, 4) ‘ubiquitin-dependent protein catabolic process’, 5) ‘translational termination’ (Figure 5). The 13 genes that correlate with cold tolerance (ENV4) show the same set of significantly enriched GO terms and in addition the GO-term ‘proteolysis’ is significantly enriched. The most significantly enriched GO-terms of the 31 genes that are associated with precipitation (ENV2) are 1) ‘regulation of pH’ and 2) ‘sodium ion transport’, followed by 3) ‘small GTPase mediated signal transduction’, 4) ‘transmembrane receptor protein tyrosine kinase signaling pathway’, 5) ‘protein phosphorylation’, and 6) ‘proteolysis’. The first two GO-terms are only associated with precipitation, while all other GO-terms are associated with at least two environmental variables (Figure 5).

Two SNPs located in EAP-OW genes associated with altitude, twelve SNPs in EAP-OW genes associated with precipitation and three SNPs in EAP-OW associated with cold tolerance are non-synonymous (Additional file 1 Figure 11) and thus we further assessed their functions (Table 3).

Amongst those EAP-OW SNPs, twelve genes are associated to different ENVs including three uncharacterized genes. The ‘proto-oncogene tyrosine-protein kinase ROS’ and the ‘breast cancer anti-estrogen resistance protein 3’ are associated to all ENVs. Both of these two genes are linked to significantly enriched GO-terms, which are enriched in the same ENVs. All other EAP-OW genes are associated to precipitation (ENV2), and one uncharacterized gene (LOC5574261) is additionally associated with cold tolerance (ENV4; Table 3). The functions of the nine characterized genes containing an EAP-OW SNP can be separated into 1) immune response, 2) life-cycle (development, reproduction, blood feeding), 3) insecticide resistance and 4) protein regulation (all details: Table 4, Figure 5).

The characteristics (such as: polar, non-polar, basic, acidic) of the amino acids before and after the base exchange demonstrate differences in the: ‘adenylate cyclase type 9’, ‘toll-like receptor Tollo’, ‘coatomer subunit beta’’ and also in two uncharacterized proteins (LOC5566519 and LOC5574261; Additional file 1 Table 7. These changes more likely can lead to a change in the protein structure or function. For all other amino acid exchanges in candidate genes, the characteristic of the amino acid stays the same.

The functional analysis of EAP-OW genes containing synonymous mutations reveals one EAP-OW gene that is associated with altitude (ENV1) playing a role in the immune response, five EAP-OW genes that are associated with precipitation (ENV2) playing a role in life-cycle (3x development, 1x blood feeding, 1x reproduction) in *Ae. aegypti* and three EAP-OW genes that are associated with cold tolerance (ENV4) involved in life-cycle (1x development, 1x reproduction) and immune response (Table 4, Additional file 2 Table 5-7). The gene ‘coatomer subunit beta’ contains two SNPs, of which one is associated with cold tolerance (ENV4) and constitutes a synonymous mutation. The other SNP constitutes a non-synonymous mutation and is associated with precipitation (ENV2).

### Genomic signatures of local adaptation

By overlapping the OW and OP window (OW-OP), 1171-1400 SNPs in 263-286 candidate genes as signatures of local adaptation are identified per population comparison (Table 2). There is no overlap between candidate genes for ‘local environmental adaptatio’ identified by Bennett (27) and candidate genes for climate adaptation (EAP-OW or EAP), Bennett (27) investigated local adaptation of *Ae. aegypti* in Panama using amongst others meteorological data of weather stations respectively. However, two candidate genes for local adaptation (OW-OP) show an overlap with the candidate genes of Bennett (27) which, however, were identified with different methods (Additional file 2 Table 8). The first gene (AAEL007657 – ‘putative vitellogenin receptor’) significantly differs between the DH600 populations and all other populations, whereas the second (AAEL002683 – ‘xanthine dehydrogenase’) significantly differs only between the CH200 and DH600 populations.

Along the altitudinal temperature gradient, knockdown resistance (*kdr*) mutations slighlty differ between populations. CH200 and KT1300 are the biggest urban sites, while CH200 and DH600 were highly effected by DENV in the last years. Thus, insecticide resistance due to a regularly insecticide use at this sites could potentially be expected. *Aedes aegypti* populations carry *kdr* mutations majorly in the biggest urban sites, respectively KT1300 followed by CH200. The V1016G mutations differ the most between populations with the wildtype (GGA) most prominently in CH200 (0.32) and KT1300 (0.44). The F1534C mutation (TGC) is major in KT1300 (0.31) compared to all other populations and no difference between populations is present in the S989P mutation (Table 5, Figure 3). None of the *kdr* mutations overlap with a significant OW/OP. Accordingly, they do not contribute to patterns of population differentiation. For the Bayesian approach, we excluded the S989P mutation, since no difference between populations was present. The Bayesian approach for comparison of the allelic combinations F1534C and V1016G points out that there is no effect of altitude on the respective allele frequencies (Additional file 1 Figure 12).

In total, 200 significant SNPs in 53 detoxification genes are associated to local adaptation, which equals to ∼0.36% of all protein-coding genes and ∼4.4% of protein coding genes involved in local adaptation (Figure 3, Figure 6, Additional File 2 Table 9). These SNPs significantly differ between populations within the OP as well as the OW (Figure 3). Out of the 200 SNPs, 113 SNPs in 30 genes are a non-synonymous mutation. The allele frequency distribution at these candidate loci were compared in a heat map revealing a slightly different pattern of frequency distribution in the KT1300 population (Figure 6). An opposite trend of allele frequency distributions is present between the KT1300 population and CH200 population as well as DK800.

In total, five SNPs in four genes are involved in vector competence, signify local adaptation, which equals to ∼ 0.03% of all protein coding genes and ∼0.3% of protein coding genes involved in local adaptation (Figure 7). Three SNPs in two genes (‘protein scarlet’, ‘leucine-rich repeat-containing protein 40’) overlap with OW-OP and have been earlier associated with DENV-1 infection by Dickson (49). Two of these three SNPs are non-synonymous SNPs. The OW-OP overlapping SNPs that are associated with DENV-3 infection in the two genes ‘cadherin-86C’ and ‘integrin alpha-PS1’ are synonymous SNPs (Figure 7, Figure 3).

**Figure 7.**
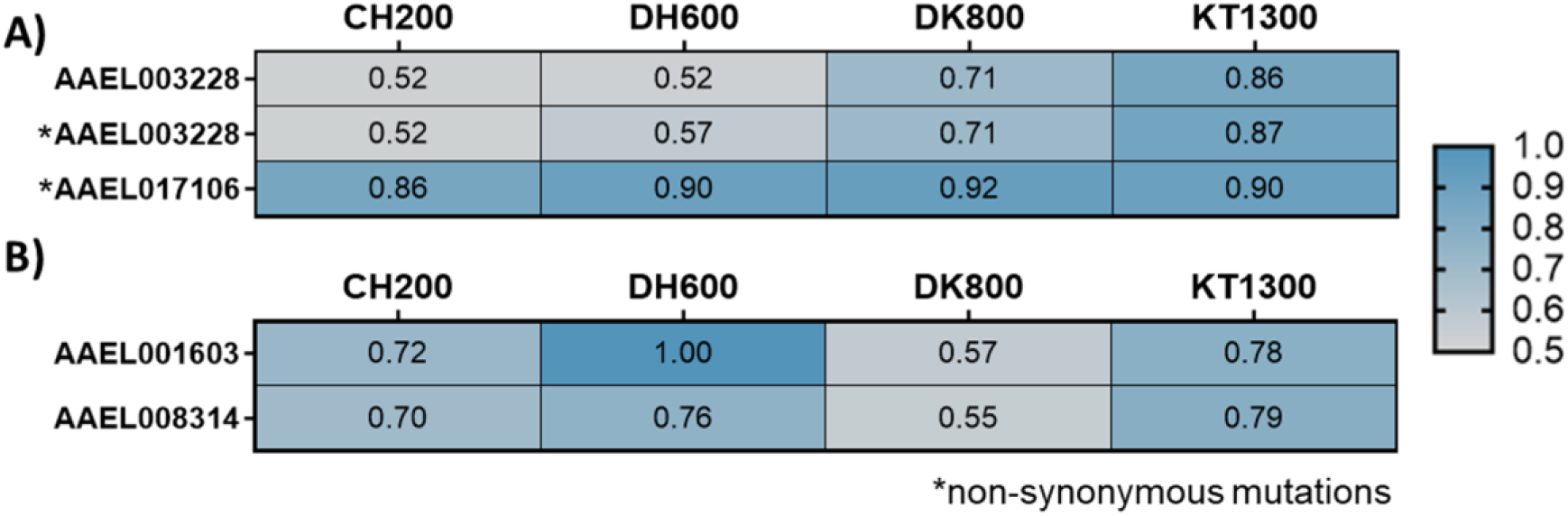
Heat map of allele frequency distribution at candidate loci associated with DENV infection. Non-synonymous (marked with a *) and synonymous mutations associated with A) DENV-1 infection or B) DENV-3 infection of *Ae. aegypti*. Altitude of sampling sites of *Ae. aegypti* populations: CH200 = 200 m asl, DH600 = 600 m asl, DK800 = 800 m asl, KT1300 = 1300 m asl.

## Discussion

The present study disentangles the genomic signature of local and climate adaption in *Ae. aegypti* populations that have been collected from an altitudinal gradient with ongoing mosquito and disease expansion to higher altitudes in the Hindukush Himalayan region (33,35,36,39,50). The observed pattern of genomic differentiation of *Ae. aegypti* populations is strongly associated to climatic differences between sampling sites. Major differences in allele frequencies uncovered 33 candidate genes for climate adaptation as well as 1200 candidate genes for local adaption. Our results specifically highlight the differing climate adaptation in the *Ae. aegypti* population sampled from the highest altitude (1300 m asl, Kathmandu) compared to the lowland populations (≤ 800 m asl) in Central Nepal. This genomic profiling of climate adaptation in *Ae. aegypti* along an altitudinal gradient contradicts our original hypothesis of a gradual expansion process of the disease vector.

### Nepalese *Ae. aegypti* populations belong to one subspecies

In comparison to worldwide *Ae. aegypti* populations, we show that all examined Nepalese populations belong to one subspecies which is most probably *Ae. aegypti aegypti* (Figure 2). This distinction was mandatory to verify that allele frequency differences were analyzed on the population but not the interspecific level. In general, it is important to distinguish the subspecies due to their likely difference in vector competence (51), even though these interspecific effects seem to depend on virus genotypes (52) and environmental factors (53, 54). Additionally, it is important to differentiate between the subspecies because of their different host preference for humans or animals (55).

### Patterns of genomic differentiation imply isolation of populations by environment

Other than the expected pattern of gradual variation along the altitudinal temperature gradient, we found significant allele frequency differences at candidate loci for climate adaptation only between the Kathmandu (KT1300) population and all other lowland populations (≤ 800 m; Figure 4). Thus, lowland populations versus the highland population form two differentiated clusters. This non-gradual pattern of genomic differentiation along the altitudinal gradient can have alternative, though not necessarily mutually exclusive, reasons. Since, the capital of Nepal (Kathmandu), is the central trading point of the country, population differentiation might derive from differences in population history such as a differential invasion history of the Kathmandu population. Alternatively, with regard to the environmental conditions along the altitudinal gradient assessed in this study, the significant differentiation in climate-associated outlier loci might be indicative for local high-altitude adaptation.

Genetic differences between Kathmandu and the lowlands might be indicative for a differential invasion history of *Ae. aegypti* in Central Nepal. To better understand the invasion process, it is important to understand how the vectors get dispersed throughout the country. The active dispersal capacity of *Ae. aegypti* is low and was reported as up to 730 m in the field (56–59). Thus, the vector expands its distribution range passively. *Aedes* mosquitoes eggs get dispersed either by the transportation of eggs in used vehicle tires (60) or through hitch-hiking of adult mosquitoes via human transportation such as aircrafts and vehicles (61, 62). *Ae. aegypti* was first recorded in Southern Nepal in 2006 (37) and since then spread rapidly throughout the country following different introduction routes along the gradient (33,35,43,46). In Kathmandu, *Ae. aegypti* was recorded for the first time in 2009 (63). The sampling sites from Chitwan (CH200) to Kathmandu (KT1300) are connected via multiple introduction roads from India (or Asia). However, since Kathmandu is the capital of Nepal and the only international airport is located there, it is thus the primary destination for any long-distance transport. This might have resulted in repeated invasion events of *Ae. aegypti* from outside of Nepal into Kathmandu. Given the clustered pattern of population differentiation between lowlands and highland populations, multiple differential or repeated invasion events across the gradient are likely. However, it has to be noted that travel and transportation routes are not unidirectional in Nepal and that invasion from Kathmandu to the lowlands is also possible. A final conclusion would, however, require a genome-wide individual-based analysis of the population structure and admixture, which cannot be performed with the given dataset.

Next to invasion history, local high-altitude adaptation exclusively in the highland population without a gradual pattern along the altitudinal gradient could imply distinct differences in environmental and climate conditions in Kathmandu when compared to the lowland population sites. This can be confirmed, since the Kathmandu climate is the coldest along the gradient, but also experiencing the harshest increase in temperature due to urbanisation, a so-called heat island effect ((41,43,64–66); Additional file 1 Figure 3+4). Nevertheless, Kathmandu represents the coldest climate where sub-zero temperatures as cold as - 2°C were present during the last years ((43); unpublished data; Phuyal, Kramer et al. 2021). We can thus conclude, that the Kathmandu climate is extreme, under strongest change, and different from the climate conditions in the lowlands, thus setting differential conditions eventually driving the isolation by environment pattern (IBE) between Kathmandu and the lowlands. Since genetic differentiation of the investigated *Ae. aegypti* populations is independent of geographic distance (see Mantel’s test) but increases with environmental differences (Figure 4), we conclude IBE over isolation by distance (IBD; (67)). Moreover, of all EAP-OWs significant SNPs were the lowest for ENV1 ∼ altitude, indicating that differences between populations is not majorly described by altitudinal geographic differences. Thus, these are optimum conditions for the identification of signatures of local adaptation without confounding demographic effects (68, 69). While the evolution of IBD is related to the interplay of genetic drift and movement, IBE is usually related to the adaptability to environmental selection pressures (70, 71). Extreme and distinct environmental and climate conditions in Kathmandu, thus, are likely to exert strong selection pressure on the highland population. The ecologically driven high-altitude adaptation is likely priming the Kathmandu population for further successful expansion into cooler habitats. After the successful establishment of populations, populations promote the speed up of the invasion by generating new introduction routes into the invaded range, the so-called bridgehead effect (summarized by (29)). In Nepal, *Ae. aegypti* is present up to 2100m altitude above mean sea level but far less abundant at altitudes above 1300m (33,35,43,46). It is unclear, if individuals present above 1300m are newly introduced each year or permanently established within the region. Thus, the established Kathmandu population can be defined as range-edge population along the investigated gradient.

The non-gradual pattern of genomic differentiation across Nepal reveals that *Ae. aegypti* bears high potential for the invasion of cooler habitats for different, mutually not exclusive reasons. Strong environmental filtering and selection is promoting high-altitude adaptation (see next section) in a population that has either been carrying a pre-adaptation due to the introduction via alternative invasion events compared to populations in the lowlands or been reaching the range-edge. The observed genomic differentiation may eventually lead to the formation of two *Ae. aegypti* lineages in Nepal, with temperate *Ae. aegypti* populations evolving along the altitudinal, as well as latitudinal gradient and a highland population with further cold tolerance adaptation. Thus, the cold tolerance and hence the fitness advantage of the high altitude population in Nepal (details on the cold tolerance potential of the Nepalese populations: (42)) may further increase (27), as also indicated by the establishment of a more cold resistant population of *Ae. aegypti* in a temperate region in Argentina (Buenos Aires; (18,72,73)). Such a phenotype would increase the introduction risk of *Ae. aegypti* into new, previously too cold ecoregions with dengue naïve human population as a process fueled by climate warming. Follow-up studies will be needed to disentangle the effects of the alternative hypotheses, ideally also investigating if individuals present at altitudes higher than Kathmandu already established and adapted to the colder climate.

### Signatures of climate adaptation in *Ae. aegypti* are genomically wide-spread and involve few genes

Here, the genomic footprint of climate adaptation could be uncovered in *Ae. aegypti*. Similar investigations were performed in different insect species, e.g. the harlequin fly (30) and two cryptic ant species (69). The investigated *Ae. aegypti* populations across the Nepalese altitudinal gradient reveal 33 candidate genes that are genomically wide-spread with signatures of climate selection, which equals to ∼0.2% of protein-coding genes. The genomic footprint of climate adaptation (i.e. adaptation to temperature and precipitation) in the harlequin fly *Chironomus riparius* involves 1.2% of protein-coding genes (30). This variation might be explained by differences in sampling design, as the altitudinal sampling gradient in Central Nepal comprised small to intermediate geographic distance, whereas Waldvogel and colleagues (30) sampled the fly populations at larger (>200 km) distances across a continental climate gradient. The here presented short-distance sampling design along a well-defined climate gradient reduces the likelihood of false-positive signals of undetected environmental variables if compared to larger scale designs incorporating higher cross-correlating heterogeneity.

Among the candidate genes of climate adaptation, significantly enriched biological processes (GO terms) either encompass general functions that are enriched to more than one environmental variables (e.g. ‘protein phosphorylation’, see Additional file 1 Table 8 for comprehensive results) or are either function specific and associated with precipitation only. As an example, the GO term ‘regulation of pH’, is associated with precipitation and is known to play a role in the hatching of larvae (74). Since for the hatching of eggs pools of rainwater are needed, the association with precipitation adds up (75).

For some of candidate genes (EAP-OW), it was possible to identify non-synonymous SNPs. Non-synonymous mutations may be associated with functional protein differences of phenotypic effect (76). We identify twelve candidate genes (EAP-OW) for climate adaptation containing non-synonymous mutations (Table 3, Table 4, Figure 5; Additional file 1 Information 1), such as the ‘toll-like receptor Tollo’. This gene was already studied in *Ae. aegypti* and plays a role in the immune response, and particularly in the anti-dengue defense (e.g. (77); details in Table 4). In addition, the non-synonymous mutations within this candidate gene lead to an amino acid with different characteristics (Additional file 1 Table 7). Since synonymous mutations may influence splicing, RNA stability, RNA folding, translation or co-translational protein folding, candidate genes (EAP-OW) containing synonymous mutations were also checked for their biological function ((78); for details Additional file 2 Table 5-7). The ‘segmentation protein Fushi tarazu’ and ‘Nasrat’ are important genes in the egg stage. The first one is involved in the segmentation in the early embryo of *Drosophila* and expresses lethal effects in *Ae. aegypti* when overexpressed, whereas the second is involved in eggshell melanization and egg viability (79, 80). These genes are 1) involved in the survival and later successful hatching of eggs and 2) associated with precipitation. The association with precipitation adds up since precipitation has an impact on survival and later successful hatching of eggs. Noteworthy, the ‘segmentation protein Fushi tarazu’ is also associated with the cold tolerance of the egg stage, indicating that cold temperature potentially affects segmentation in the embryo of *Ae. aegypti* (79). For verification, knock-out studies testing the molecular function of the *Ae. aegypti* candidate genes containing different SNPs at given positions are highly recommended.

### Signatures of local adaptation reveal a broad functional basis in *Ae. aegypti*

Other than gradual climate selection regimes, local selection pressures act on populations only in their specific habitat. Accordingly, there are SNPs that are not associated to the climatic gradients but still highly divergent between some or all *Ae. aegypti* populations (OW-OP). These SNPs are candidates for local adaptation. Approximately 8.2% of protein-coding genes, i.e. 1200 genes, show signatures of local selection. Similarly, 7.6% of genes were found to be locally adapted in *C. riparius* (30). Two of the identified candidate genes for local adaptation were already found to play a role in local adaptation of *Ae. aegypti* in Panama (27). Due to the identification of these two genes in *Ae. aegypti* populations from different countries, the two genes seem to play an important role in local adaptation of this species. The first gene ‘putative vitellogenin receptor’ significantly differs between the DH600 population *versus* the other populations and the second ’xanthine dehydrogenase’ only significantly differs between the CH200 and DH600 population. The tropical climate at the respective lowland populations (CH200, DH600) and the populations from Panama support the indication that the genes could be important in coping with tropical climate variables such as high humidity or high temperature. In general, it is known that the ‘putative vitellogenin receptor’ plays a role in the vitellogenesis (yolk formation) of *Ae. aegypti* females and is increasingly upregulated post-emergence prior to the first gonotrophic cycle (81) while the ‘xanthine dehydrogenase’ is involved in survival of blood-fed *Ae. aegypti* mosquitoes. Silencing of this gene influences digestion, excretion and reproduction. Due to the lethal effect in blood-fed mosquitoes, this gene could be targeted to control vector populations (82).

Amongst all candidate genes for local adaptation, we spotlight two traits that are important from a medical vector-borne disease perspective, namely insecticide resistance and vector competence. The insecticide resistance of *Ae. aegypti* determines the success of vector control programs (47). Most variations with the detoxification enzymes are probably not functionally associated with insecticide resistance. Instead some are the consequence of strong selection pressure, hence only some reflect selection of a variant showing an increased metabolic activity against insecticides (47). However, *kdr* mutations such as V1016G, F1534C and S989P are known to lead to pyrethroid insecticide resistance in *Ae. aegypti* (summary in (47)). In accordance with Kawada (82), in Nepal the *kdr* mutations F1534C and V1016G are present with varying frequencies and the S989P mutation was not present in all study populations (Table 5, Figure 3). Within the CH200 and KT1300 population, there is a trend of increased *kdr* mutations. It can be hypothesized that this trend is present since fogging of insecticides (deltamethrin) mainly occurs in urban areas. Thus *kdr* mutations may be more present in urban areas such as CH200 and KT1300 compared to less urban regions such as DK800 and DH600 (82). Kawada (82) showed at least for CH200 and KT1300 their susceptibility to pyrethroids. However, none of the *kdr* mutations are found to overlap with a significant OW/OP and accordingly they did not contribute to patterns of population differentiation. Given that the Nepalese populations showed an intermediate to high resistance to pyrethroids, but only small amounts of insecticides are used in Nepal compared to other Asian countries (82), this indicates a reduced selection pressure on *kdr* mutations in Nepal. The genetic presence of *kdr* mutations might derive from the introduction of *Ae. aegypti* populations from neighboring countries (82), most likely from India.

The vector competence of *Ae. aegypti* determines the efficiency of dengue transmission to humans and thus it is important to understand this trait at a local level. SNPs associated with DENV-1 and/or DENV- 3 infection were found in all populations and likely play a role in dengue resistance of *Ae. aegypti* in Central Nepal. This assumption is supported by reported DENV type-specific resistance of a population from Gabon (49). Interestingly, the candidate gene ‘integrin alpha-PS1’ has already been proven to play a role in infection of bluetongue virus in *Ae. albopictus* (83). The synonymous mutation in this candidate genes ‘integrin alpha-PS1’ (EAP-OW) may influence splicing, RNA stability, RNA folding, translation or co-translational protein folding (78), since in infected *Ae. albopictus* cells, mRNA of the candidate gene was upregulated. One may speculate, that the candidate gene ‘integrin alpha-PS1’ influences the dengue virus dissemination, replication and transmission efficiency in *Ae. aegypti*.

However, the verification of SNPs and their functional meaning in the identified candidate genes for insecticide resistance and dengue vector competence merits definitely further research.

## Conclusion and implications for climate adaptation

In a worldwide comparison with other *Ae. aegypti* populations we showed that Nepalese mosquitoes belong to a single subspecies. Patterns of genomic differentiation between the 1300 m population in comparison to all other lowland populations (≤ 800 m) imply isolation by environment (IBE). By demonstrating a distinct genomic footprint of climate adaptation in *Ae. aegypti*, our study assists to close the knowledge gap on adaptive traits and associated gene sets on climate adaptation of *Ae. aegypti* (31), while signatures of local adaptation reveal a broad functional basis of the species. In total, twelve candidate genes (EAP-OW) for climate adaptation containing non-synonymous mutations were identified. Amongst all candidate genes for local adaptation, we spotlight two traits important from a medical VBDs perspective, namely insecticide resistance and vector competence.

Genomic differentiation of the 1300 m population compared to the lowland populations either indicate invasion of a pre-adapted population due to an alternative invasion route compared to the lowland populations or local adaptation of the 1300 m range-edge population. In any case, the identified alleles of the highland population are likely relevant for their invasion to colder regions. In general, it is of major importance to track the trends of climate adaptation not only in emerging viruses (84), but also in the respective vector populations especially. On the most basic level, differentially adapted populations, be it to climate or local conditions, could have different abilities to transmit arbovirus diseases (27). With our study we demonstrate that effective monitoring of vector populations using NGS strategies allows to interpret emerging expansion trends, and especially population samples proved to be a powerful and cost-effective methodology to assist the comprehensive monitoring and mapping of the vector species *Ae. aegypti* (85). Patterns of population differentiation, genomically as well as physiologically, deliver important evolutionary and ecological information to be integrated into vector distribution models or VBDs risk assessments under climate change scenarios, especially in cooler ecoregions (44). Thus, current distribution models predicting the future distribution of vector populations should incorporate the adaptive response of species for more precise predictions (27). Genomic diversity and thus biodiversity by means of adaptation and simultaneously climate warming is likely to increase the risk of expansion of *Ae. aegypti* worldwide to colder ecoregions. With the increasing distribution range of the vectors worldwide as well as in Nepal and the HKH region in particular (9,19,20) also the spread of VBDs will increase (worldwide: e.g. dengue: (6); Nepal: (86)), underlining that parts of biodiversity can be detrimental to human health. For efficient vector control, it is important to consider that locally adapted populations could impact control efforts that are based on gene drive system, but adaptive genes could also be targets for population control using gene editing strategies (87–89).

Results obtained in this study could potentially be used for the inference of the adaptive response of *Ae. aegypti* to colder ecoregions worldwide. The health systems in cooler ecoregions need to prepare for future VBD outbreaks and develop surveillance strategies to prevent the establishment of dengue vectors. To identify emerging trends within the adaptation of *Ae. aegypti* to new environments, we recommend to investigate populations in Nepal from higher altitude as well as populations along altitudinal and latitudinal clines worldwide. Moreover, next to reciprocal transplant experiments (27, 87), molecular investigations of the function of the candidate genes, the verification of the association of candidate genes with different environmental variables and differences in vector competence between the KT1300 populations and lowland populations should be verified.

## Methods

### Collection of mosquitos

We sampled *Ae. aegypti* populations, each with a minimum of 100 individuals, from four sampling sites: Chitwan (CH200, 200 m above sea level), Dhading (DH600, 600 m asl), Dharke (DK800, 800 m asl) and Kathmandu (KT1300, 1300 m asl). The sampling sites are distributed along an altitudinal and temperature gradient in Central Nepal ((42, 46); Figure 1) and connected via a motorway (Chitwan → Dhading (side valley; road distance:∼97 km) → Dharke (∼57 km) → Kathmandu (∼31 km)). *Aedes* larvae, pupae and adults that were available in/near temporary water reservoirs, such as containers or tires, were collected during the high mosquito season (late monsoon and early post-monsoon; September till October 2018; (46). Immature stages were reared to adults using paper cups covered with a net and water from their respective sampling site. If less than 100 *Ae. aegypti* individuals (larvae, pupae, adults) were sampled in the field, eggs from the same sampling campaign were reared to adulthood at the Department of Environmental Toxicology & Medical Entomology, Institute of Occupational, Social and Environmental Medicine; Goethe University Frankfurt, Germany (more details in Additional file 1 Table 1 and (42, 46). either sampled or emerged from rearing were conserved in 100% ethanol. Dead mosquitoes were identified by a local taxonomist following the guidelines described in (32). This combination ensured that all individuals of the pool represented true field samples, only differing in the developmental stages at the time point of sampling. For DNA isolation (Qiagen DNeasy Blood and Tissue kit, Hilden, Germany), two legs of each adult mosquito were pooled per population. To control the quantity of DNA, Qubit® Fluorometer (Invitrogen, Massachusetts- USA) measurements were performed.

### Pool-Seq genome scans

Four pooled DNA samples were sequenced on an Illumina HiSeq to yield 150 bp paired-end pooled sequencing (Pool-Seq) whole genome data (Figure 1). The ratio of ≥96 individuals per population and targeted coverage of ∼20-30X per pool was chosen to allow an accurate estimation of genome-wide allele frequencies (90, 91). Pool-Seq genome data were quality trimmed and separately pre-processed using the wrapper script *autotrim.pl* ((30), available at https://github.com/schellt/autotrim), which integrates *Trimmomatic* (92) and *fastqc* (93).

### Analysis of subspecies: Microsatellite analysis

To link our population genomic analyses to prior microsatellite work and to identify potential subspecies as they are described for *Ae. aegypti* (94), we developed a workflow to assess microsatellite (µsats) diversity from genome-wide Pool-Seq data. For this analysis explicitly, the trimmed files were mapped to the unmasked reference genome of *Ae. aegypti* (48) using *NextGenMap* (*ngm*, (95)). Accounting for the possible presence of subspecies of *Ae. aegypti* (dominant African subspecies: *Ae. aegypti formosus*; outside of Africa: *Ae. aegypti aegypti* (94)) in the samples, *ngm* was used since this mapper is independent of the amount of genomic polymorphism present in reads (95). Each read of genome-wide Pool-Seq data belonging to one individual chromosome (diploid individuals), provides the required haplotype-specific data to analyse population structure using µsats. First, 12 µsats were identified (A1, A9, AC1, AC2, AC4, AC5, AG2, AG4, B2, B3, CT2, AG1; (94)), located and extracted along the reference genome via the *in_silico_PCR.pl* script (https://github.com/egonozer/in_silico_pcr) and making use of primers from Brown and Slotman (96, 97). AG1 could not be identified along the reference genome and was therefore excluded from the analysis. Following the identified coordinates of the reference genome, µsats alignments were extracted from mapped bam files using *samtools* (98). Each µsat alignment was re-aligned to the extracted µsat reference sequence and, if available, to the Slotman (97) reference sequences of the respective µsats. Alignments were manually edited using Geneious Prime® 2019.2.1. Repeated elements were identified either using the µsats reference (97) or the MISA-web tool (99). As a measure of quality filtering, re-aligned sequences (single sequences = haplotypes) were included only if each µsat covered at least 2 bp before the start and behind the end of satellite region. Gaps and duplicates were removed and start and end positions of sequences were set to Ns to fix the alignment structure when saving the data in fasta format. Counting repeated elements (in bp) per µsats and individual, their frequencies per population were calculated. Using this population frequency data, 50 individuals were simulated with a custom Python script under the assumption of Hardy-Weinberg equilibrium in order to make our data comparable to individual frequency data. Individuals were only simulated if a minimum amount of four reads was present at a µsat.

To compare this data with a world-wide set of populations and to test for the presence/absence of subspecies in Nepal, the same workflow was followed using publically available genome-wide data of four laboratory populations (West Africa – likely from Freetown-Sierra Leone belonging likely to *Ae. aegypti formosus*, likely *Ae. aegypti aegypti*: Australia – Innisfail, USA – Clovis, Costa Rica – Puntarenas) comprising each 30 females (individual sequencing; (48, 101); Accession number: SRX3413563-SRX3413566). Only µsats with a coverage higher than or equal to four individuals were used for the analysis of population structure (used µsats: A9, AC1, AC4, AG2, B2, B3; Additional file 1 Table 2). The population from the USA was excluded due to low individual coverage of this specific data set (Additional file 1 Table 2). Using the Bayesian clustering method implemented in the software STRUCTURE v. 2.3.4 (101), the population structure as described in (94) was assessed. Each conducted run assumed an admixture model and correlated allele frequencies with a burn-in of 250,000 iterations with in addition 750,000 repetitions. To specifically test for differences between all populations and the African population, the structure analysis was performed with K=2 (compare with (94)) with ten iterations. To summarize STRUCTURE results of the ten iterations per K and plot consistent cluster coloring CLUMPAK was used (102). In order to exclusively assess differences among the populations of Nepal the population structure with K=1-4 was calculated.

### Genome wide population differentiation

Estimation of population differentiation using the genome wide SNP data followed the pipeline of *PoPoolation2* (103) and (30). Before mapping, overlapping read pairs were assembled using *PEAR* (104). This was necessary in order to make use of the full data set, though only a small proportion of reads were found to overlap, while avoiding erroneous allele frequency estimates in overlapping regions. Assembled and unassembled reads were mapped to the available reference genome (masked version) of *Ae. aegypti* (48) sing *bwa mem* (105). Duplicates were removed using *picard tools* (106) and all bases below a minimum mapping quality of 10 were discarded (*samtools;* (98))*. PoPoolation* (107) was used to estimate population specific parameters such as the nucleotide diversity (π; genome-wide per site and in 1kb window, exon-wide per site) and the population mutation parameter (θ; genome-wide in 1kb window). The effective population size (Ne) was calculated using genome-wide θ estimates as follows: 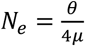. The genome wide mutation rate (μ) of *Chironomus riparius* was used for the N_e_ calculation (108).

For comparative analyses between populations, the pipeline *PoPoolation2* was followed (103). In brief, pairwise FST values (*fst-sliding.pl*) of all population pairs in a sliding window of 1kb along the subsampled *sync*-file were calculated. The upper 1% tail of the FST distribution was defined as threshold for non-neutral differentiation, as this has been shown to provide a conservative threshold for a robust drift expectation (30). In addition, for each 1 kb-window Fisher’s p-values (*fisher-test.pl*) were calculated and the Benjamini-Hochberg correction against multiple testing to all p-values was performed. We defined highly significant outlier windows (OW) to be those windows that remained significant after FDR correction (q < 0.01). Circos tool was used to graphically illustrate the distribution of OWs along the genome (109). As described for the OW estimation we additionally calculated highly significant outlier positions for each population (OP) per site. In addition, to test for genome-wide isolation by distance patterns, a Mantel test with 23 permutations (complete enumeration) in R/VEGAN (110) between the genome-wide mean FST values and the geographical distance was calculated.

### Environmental data

The following environmental data of *Aedes* sampling sites were analysed to provide the environmental data for the GEA: i) microclimate data/logger data (temperature data; Additional file 1 Table 3 and 4), ii) high-resolution data from CHELSA of 1979–2013 (Additional file 1 Table 3), and iii) Bioclim variables ((111); data source: (112); 30 arcsec, ∼1 km from CHELSA version 1.2; Additional file 1 Table 3). HOBO data loggers (type UX100-011A, ONSET®) were installed indoors in houses with no heating or air condition and bad isolation (I) and outdoors at shaded artificial places (SH; e.g. near households) at sampling sites from November 2017 to March 2019. Loggers were additionally installed at 1800 and 2050 m asl (Ranipauwa =RP1800, 1800 m asl; Dhunche= DU2050, 2050 m asl). In Dharke (800 m), HOBO loggers were missing and thus the data of the 800 m sampling site were interpolated from logger data obtained along the altitudinal gradient of 200 m to 2050 m asl using linear regression (Prism®, Version 7, GraphPad Software Inc., USA). By means of a principal component analysis (PCA), confounding covariation in the environmental data set was reduced.

### GEA

To analyse how the genomic differentiation is potentially correlated with environmental variation across sampling sites, a genotype-environment association analysis was performed using LFMM (Latent Factor Mixed Model) in the frame of the ‘LEA’ R-package (113), which is amongst the most commonly applied tools in GEA studies (45). The Pool-Seq approach does not account for pool size (30) and thus 20 pseudo-individual allele frequency spectra were inferred by simulating observed allele frequencies at each locus referring to the BAYENV approach (114). In accordance, for each locus environmental factors were replicated 20 times. Considering the large genome size of *Ae. aegypti* as well as the main target to identify candidate genes in downstream analysis only the coding regions (CDS) were included. Three PCA components and the cold tolerance (normalized mean survivorship after cold exposure to - 2°C for 8 days to controls; CT; ENV4; (42)) were used as environmental input variables (ENV) for the GEA (Additional file 1 Table 6). We ran the LFMM function “lfmm_ridge” with a latent factor of K = 4 (reflecting number of populations; algorithm = analytical). p-values were calibrated by computing the median and MAD (Median Absolute Deviation) of the *z* scores using the “lfmm_test” function (Additional file 1 Figure 8, Additional file 1 Table 6). We ran LFMM twice for different combinations of environmental input variables: 1) PCA1 – altitude, 2) PCA2 – precipitation, PCA3 – seasonality plus PCA4 – CT (Additional file 1 Table 5, Additional file 1 Figure 6-9). Resulting output p-values were FDR corrected and positions with q < 0.01 defined as significant ENV associated positions (EAP).

We then compared the highly differentiated outlier windows (OW) from our previous analysis on population differentiation, with the here resulting set of EAPs and checked for overlapping regions of the two sets. With regard to the above-described characteristics to define OWs, we again stringently considered only those EAPs, which fell into a respective OW (EAP-OW). Differences in allele frequencies of the candidate SNPs (EAP-OW positions) along the gradient were analyzed per ENV using Prism® (Version 7, GraphPad Software Inc., USA). In order to identify highly significant positions for climate adaptation, we verified if EAP-OW are additionally present in those OPs.

### Functional enrichment associated with climate adaptation

Candidate genes were studied in a functional enrichment analysis. Therefore, all EAP-OW positions were annotated using the coordinates of protein coding genes of the *Ae. aegypti* reference genome (48). InterProscan (115) was used to classify proteins into families and predicting domains as reference for the functional enrichment analysis. Gene ontology (GO) terms significantly enriched in genes were then analyzed using the topGO R package (116) in the category ‘biological processes’, with the weight01 algorithm and Fisher statistics. Enriched GO terms with a p < 0.05 were further assessed (30). To analyze if base substitutions at SNPs lead to synonymous or non-synonymous mutations in the amino acid sequence of candidate genes, tbg-tools v0.2 (https://github.com/Croxa/tbg-tools; (76)) was used. The characteristic of the amino acid present, before and after the base exchange was also assessed (117, 118).

Knowledge on the biological function of candidate genes containing non-synonymous mutations was collected from literature and databases. We performed a literature survey in Google Scholar by using the candidate gene name (or/and the Locus tag) in combination with the following terms: 1) *Aedes aegypti*, 2) *Aedes,* 3) mosquito, 4) insect. Furthermore, we extracted candidate gene IDs containing non-synonymous and synonymous mutations and searched for their function using UniProt, NCBI, and Vectorbase. Moreover, we screened GO-terms of candidate genes in UniProt for similarities of GO-terms found in the functional enrichment analysis. The procedure was likewise repeated also for candidate genes containing synonymous mutations but only the locus tag and the species name was used as a search term.

### Genomic signatures of local adaptation

Next to climate adaptation, we searched for candidate genes indicating strong local adaptation. Therefore, we defined candidate genes/positions laying in the CDS, that were not overlapping with an EAP but were present in an OW and additionally overlapped with an OP (OW-OP), as candidates for purely local adaptation. Potential candidate genes for local adaption that are involved in insecticide resistance or vector competence were especially taken into focus. Additionally, we compared candidate SNPs/genes of the Nepalese population with a recent study that investigated genomic signs of ‘local environmental adaptation’ (=climate adaptation) in populations from Panama (17 genes; (27)).

We located the *kdr* mutations V1016G, F1534C, and S989P in the reference genome and extracted the sequences from one sorted *bam-*file of one population by using the *in_silico_PCR.pl* script (https://github.com/egonozer/in_silico_pcr) and the primers given by (119). Extracted sequences were processed in Geneious Prime® 2019.2.1 and excact genome positions of the *kdr* mutations were calculated. Allele frequencies at the position of the *kdr* mutations were extracted from the *sync*-file and overlaps of *kdr* mutations with OW as well as OP were checked.

The combined occurrence of the *kdr* mutations in the populations sampled along the altitudinal gradient was investigated using allele frequency differences. We fitted a Bayesian multivariate response model with binomial distribution of the allele frequency differences of *kdr* mutations with the brms package (120), which is a high-level interface to Stan (121) with R v.4.0.5 (122) in RStudio v.1.3.959 (123). The response variable “allele frequency” was included as the proportion of the major allele observations to all allele counts using “trials”. In addition to the fixed factor “altitude”, an “additive overdispersion” random effect was added to estimate the residual correlation. The model was run without intercept, and additionally without the residual random effect as well as without altitude and tested for differences between those models using the “leave-one-out” criterion. As the model fit did not differ between models, the full model including altitude and the random factor is reported only. The full model was run with 4 parallel chains with 3,500 iterations each, where the first 1,000 were used as warm up and discarded. Priors were flat for allele frequencies as suggested by the “get_prior” function. Trace plots, effective sample sizes (range of effective sample sizes: 755 – 4822) and R-hat (124) values (1 < 1.02) confirmed a proper convergence.

Allele frequency differences of detoxification genes (as listed by (47)) were checked in our dataset and in the genome annotation published by (48) for i) being part of the CDS, ii) having an overlap with OW- OP and iii) showing a non-synonymous or synonymous mutation (tbg-tools v0.2; https://github.com/Croxa/tbg-tools; (76)). Differences in allele frequencies at candidate SNPs between populations were visualized in a heat map (Prism®, Version 7, GraphPad Software Inc., USA).

Local adaptation in vector competence was analyzed by comparing a list of SNPs (top 0.001% most significant SNPs) associated with DENV1 or/and DENV3 infection by (49) with the allele frequency at the respective site of the Nepalese populations. With a stringent approach, we checked whether these SNPs were present in the CDS, overlap with OW-OP and whether SNPs lead to a non-synonymous or synonymous mutation with the tbg tool. As described above, the allele frequencies at candidate SNPs were visualized using a heat map to compare them between the Nepalese populations, to identify different resistance to dengue infection (Prism®, Version 7, GraphPad Software Inc., USA).

## Declarations

### Ethics approval and consent to participate

The conduct of this study was approved by the Ethical Review Board of the Nepal Health Research Council (NHRC), Government of Nepal (381/2017).

### Consent for publication

Not applicable

### Availability of data and materials

The PoolSeq-datasets supporting the conclusions of this article are currently uploaded to ENA- European Nucleotide Archive. In the next version accession numbers will be added.

### Competing interests

The authors declare that they have no competing interests.

### Funding

The work was funded by the Federal Ministry of Education and Research of Germany (BMBF) under the project AECO (Number 01Kl1717) as part of the National Research Network on Zoonotic Infectious Diseases of Germany. This work was additionally supported by the LOEWE Centre Translational Biodiversity Genomics (TBG), funded by the state of Hesse, Germany.

### Authors’ contributions

IMK, MD, IG, PS, SB and RM sampled the mosquito populations. PS entomologically identified the mosquito species. IMK and AMW majorly analysed the data. MP, BF, JH, AM, BA and RM assisted in the data analysis. IMK, BF, PP, JH, RM and AMW visualized the data. MD, RM and AMW conceptualized the study. MP, DAG and RM provided study resources. RM and AMW supervised the study. RM and AM were responsible for the study administration and RM also for the funding acquisition. IMK and AMW wrote the original draft. MP, BF, MD, IG, PS, SB, PP, JH, AM, DAG, BA and RM reviewed and edited the original draft. All authors read and approved the final manuscript.

## Acknowledgements

The authors wish to thank Sabita Oli, Keshav Luitel and Sandhya Belbase from the Nepal Health Research Council (Kathmandu, Nepal) for the help in the adult/egg sampling campaign in Nepal. We also thank Dr. Tilman Schell and Christoph Sinai from the TBG- LOEWE Centre for translational biodiversity genomics for their support for the population genomic analysis. Moreover, we thank Doris Klingelhöfer for her support in the figure design.

## Additional Files

**Additional File 1 (.txt): Table 1.** The number, sex and original life-stage of *Ae. aegypti* individuals used per Pool-Seq sample. **Table 2.** Number of individuals that cover a microsatellite region of eight populations using PoolSeq data. **Table 3.** Resolution of environmental data used for PCA. **Table 4.** Detailed description of logger data and their installation period in the field. **Table 5.** Climate variables and Bioclim dataset used in the PCA. **Table 6**. LFMM median values per sampling site and environmental variables. **Table 7**. Characteristic of amino acid before and after alternative base exchange at non-synonymous SNP position. **Table 8.** Significantly enriched GO terms among candidate genes and their biological functions involved in climate adaptation. **Figure 1**. Delta K and Probability by K from the STRUCTURE analysis. **Figure 2**. Pairwise FST distribution per 1 kb-windows of Nepalese *Ae. aegypti* populations. **Figure 3**. Climate along the altitudinal gradient in Central Nepal. **Figure 4**. Microclimate along the altitudinal gradient in Central Nepal. **Figure 5**. Precipitation along the altitudinal gradient in Central Nepal. **Figure 6**. Loadings from PC (principal component) analysis: PC1 is associated with altitude. **Figure 7**. Loadings from PC (principal component) analysis: PC2 is associated with precipitation. **Figure 8.** Loadings from PC (principal component) analysis: PC3 is associated with seasonality. **Figure 9.** Distribution of Eigenvalues (%) of principal components (blue line). **Figure 10.** The frequency distribution of adjusted p-values after association to four different environmental variables using LFMM. **Figure 11.** Gene IDs or protein IDs present in all different significant environmental variable associated positions laying in an overlapping singificant 1kb-FST-window and contain a non-synonymous mutation. **Figure 12.** Posterior uncertainty intervals for *kdr* mutation. **Figure 11**. Pairwise FST distribution per 1 kb-windows of Nepalese *Ae. aegypti* populations. **Information 1**. Details on GO terms and Candidate genes containing non-synonymous mutations.

**Additional File 2 (.xls): Table 1:** General file information about the amount of candidate genes and SNPs. Table 2: Gene description of ENV1 candidate genes from the LFMM analysis (Presence of candidate SNPs in OWP or OW is indicated). Table 3: Gene description of ENV2 candidate genes from the LFMM analysis (Presence of candidate SNPs in OWP or OW is indicated). Table 4: Gene description of ENV4 candidate genes from the LFMM analysis (Presence of candidate SNPs in OWP or OW is indicated.). Table 5: Detailed gene description of highly significant candidate genes for climate adaptation associated with ENV1 laying in an OW. Table 6: Detailed gene description of highly significant candidate genes for climate adaptation associated with ENV2 laying in an OW. Table 7: Detailed gene description of highly significant candidate genes for climate adaptation associated with ENV4 laying in an OW. Table 8: Candidate genes for local adaptation (presence in EAP-OW and (27) is indicated). Table 9: List of Detoxification genes containing non-synonymous mutations

## Notes

### Competing Interest Statement

The authors have declared no competing interest.

