## Additional File 1 for "Genomic profiling of climate adaptation in *Aedes aegypti* along an altitudinal gradient in Nepal indicates non-gradual expansion of the disease vector"

**Table 1.** The number, sex and original life-stage of *Ae. aegypti* individuals used per Pool-Seq sample. The set-up of Pool-Seq samples comprising DNA of ≥96 adult individuals per *Ae. agypti* population from four sampling site in Central Nepal. Immature life-history stages were reared to adulthood prior to DNA isolation. Altitude of sampling sites of *Ae. aegypti* populations: CH200 = 200 m asl, DH600 = 600 m asl, DK800 = 800 m asl, KT1300 = 1300 m asl.

|  | **CH200** | | **DH600** | | **DK800** | | **KT1300** | |
| --- | --- | --- | --- | --- | --- | --- | --- | --- |
|  | F | M | F | M | F | M | F | M |
| **individuals per sex** | 51 | 51 | 51 | 51 | 53 | 43 | 52 | 51 |
| **larvae/pupae/adult (F0)** | 32 | 35 | 51 | 51 | 42 | 33 | 52 | 51 |
| **eggs (F0)** | 19 | 16 | - | - | 11 | 10 | - | - |
| **Individuals in total** | **102** | | **102** | | **96** | | **103** | |

**Table 2.** Number of individuals that cover a microsatellite region of eight populations using PoolSeq data. Regions with a less than 4 individuals are marked in bold. Altitude of sampling sites of *Ae. aegypti* populations: CH200 = 200 m asl, DH600 = 600 m asl, DK800 = 800 m asl, KT1300 = 1300 m asl.

| **Microsatellite** | **Liverpool**  **(West Africa)** | **Innisfail**  **(Australia)** | **Clovis**  **(USA)** | **Puntarenas**  **(Costa Rica)** | **CH200**  **(Nepal)** | **DH600**  **(Nepal)** | **DK800**  **(Nepal)** | **KT1300**  **(Nepal)** |
| --- | --- | --- | --- | --- | --- | --- | --- | --- |
| A1 | **1** | **0** | **3** | 9 | 29 | 17 | 23 | 18 |
| A9 | 12 | **3** | 5 | 21 | 22 | 18 | 25 | 29 |
| AC1 | 4 | **2** | **1** | 14 | 10 | 12 | 28 | **2** |
| AC2 | **1** | 6 | **2** | 5 | 27 | 34 | **0** | 10 |
| AC4 | 9 | 5 | 8 | 17 | 44 | 27 | 42 | 31 |
| AC5 | **3** | 3 | **1** | 7 | 20 | 10 | 58 | 26 |
| AG2 | 8 | 9 | **1** | 8 | 20 | 19 | 10 | 24 |
| AG4 | **3** | 4 | **3** | 22 | 26 | 29 | 38 | 24 |
| B2 | 4 | 5 | **3** | 12 | 21 | 27 | 27 | 21 |
| B3 | 6 | **3** | **1** | 11 | 35 | 23 | 9 | 15 |
| CT2 | **2** | 8 | 4 | 9 | 49 | 28 | 35 | 26 |

**Table 3.** Resolution of environmental data used for PCA. Microclimate has been derived from logger data and regional climate estimates from CHELSA (including all Bio variables 1-19; (1))

|  | **mean temperature** | **minimum temperature** | **maximum temperature** | **precipitation** |
| --- | --- | --- | --- | --- |
| **Logger data** | per season/annually | per season/annually | per season/annually |  |
| **CHELSA** | per month | per month | per month | per month |

**Table 4.** Detailed description of logger data and their installation period in the field. I = indoors; SH= shadowed artificial substrates. Altitude of sampling sites of *Ae. aegypti* populations: CH200 = 200 m asl, DH600 = 600 m asl, DK800 = 800 m asl, KT1300 = 1300 m asl. SH logger at DH600 was lost during data recording.

| **Altitude [m]** | **Microclimate** | **Start 2017** | **End 2019** | **Missing data** |
| --- | --- | --- | --- | --- |
| CH200 | I1 | 16.11.2017 | 03.2019 |  |
|  | I2 | 16.11.2017 | 03.2019 |  |
|  | SH | 16.11.2017 | 03.2019 |  |
| DH600 | I1 | 01.02.2018 | 03.2019 |  |
|  | I2 | 16.11.2017 | 03.2019 |  |
|  | SH | Missing logger | |  |
| DK1300 | I1 | 15.11.2017 | 03.2019 |  |
|  | I2 | 15.11.2017 | 03.2019 |  |
|  | SH | 12.11.2017 | 03.2019 |  |
| RP1800 | I1 | 03.10.2018 | 03.2019 |  |
|  | I2 | 03.10.2017 | 03.2019 |  |
|  | SH | 03.10.2017 | 03.2019 | 3.10.18-7.10.18 |
| DU2050 | I | 09.11.2017 | 03.2019 |  |
|  | I2 | 09.11.2017 | 03.2019 |  |
|  | SH | 09.11.2017 | 03.2019 |  |

**Table 5.** Climate variables and Bioclim dataset used in the PCA. The respective highest components per PC loading are given. Climate variables were collected from HOBO logger datasets 11/2017-03/2019 and CHELSA (C.) database (1979-2013) and the Bioclim dataset from 1979-2013 (see also Additional File 1 Figure 6-8). Temperature = Temp; Precipitation = Prec; Minimum = min; Maximum = max.

| **Term of Parameter** | **related to** | **Logger** | **CHELSA** | **Bioclim** | **PC1** | **PC2** | **PC3** |
| --- | --- | --- | --- | --- | --- | --- | --- |
| **Longitude** |  | X |  |  |  |  |  |
| **Latitude** |  | X |  |  |  |  |  |
| **Altitude** |  | X |  |  | 0.97 |  |  |
| **Annual mean** | Temperature | X |  |  |  |  |  |
| **Annual min** | Temperature | X |  |  |  |  |  |
| **Annual max** | Temperature | X |  |  |  |  |  |
| **Pre-monsoon mean** | Temperature | X |  |  |  |  |  |
| **Pre-monsoon min.** | Temperature | X |  |  |  |  |  |
| **Pre-monsoon max.** | Temperature | X |  |  |  |  |  |
| **Monsoon mean** | Temperature | X |  |  |  |  |  |
| **monsoon min.** | Temperature | X |  |  |  |  |  |
| **Monsoon max.** | Temperature | X |  |  |  |  |  |
| **Post-monsoon mean** | Temperature | X |  |  |  |  |  |
| **Post-monsoon min.** | Temperature | X |  |  |  |  |  |
| **Post-monsoon max.** | Temperature | X |  |  |  |  |  |
| **winter mean** | Temperature | X |  |  |  |  |  |
| **winter min.** | Temperature | X |  |  |  |  |  |
| **winter max.** | Temperature | X |  |  |  |  |  |
| **Max. January C.** | Temperature |  | X |  |  |  |  |
| **Max. February C.** | Temperature |  | X |  |  |  |  |
| **Max. March C.** | Temperature |  | X |  |  |  |  |
| **Max. April C.** | Temperature |  | X |  |  |  |  |
| **Max. May C.** | Temperature |  | X |  |  |  |  |
| **Max. June C.** | Temperature |  | X |  |  |  |  |
| **Max. July C.** | Temperature |  | X |  |  |  |  |
| **Max. August C.** | Temperature |  | X |  |  |  |  |
| **Max. September C.** | Temperature |  | X |  |  |  |  |
| **Max. October C.** | Temperature |  | X |  |  |  |  |
| **Max. November C.** | Temperature |  | X |  |  |  |  |
| **Max. December C.** | Temperature |  | X |  |  |  |  |
| **Min. January C.** | Temperature |  | X |  |  |  |  |
| **Min. February C.** | Temperature |  | X |  |  |  |  |
| **Min. March C.** | Temperature |  | X |  |  |  |  |
| **Min. April C.** | Temperature |  | X |  |  |  |  |
| **Min. May C.** | Temperature |  | X |  |  |  |  |
| **Min. June C.** | Temperature |  | X |  |  |  |  |
| **Min. July C.** | Temperature |  | X |  |  |  |  |
| **Min. August C.** | Temperature |  | X |  |  |  |  |
| **Min. September C.** | Temperature |  | X |  |  |  |  |
| **Min. October C.** | Temperature |  | X |  |  |  |  |
| **Min. November C.** | Temperature |  | X |  |  |  |  |
| **Min. December C.** | Temperature |  | X |  |  |  |  |
| **Mean January C.** | Temperature |  | X |  |  |  |  |
| **Mean February C.** | Temperature |  | X |  |  |  |  |
| **Mean March C.** | Temperature |  | X |  |  |  |  |
| **Mean April C.** | Temperature |  | X |  |  |  |  |
| **Mean May C.** | Temperature |  | X |  |  |  |  |
| **Mean June C.** | Temperature |  | X |  |  |  |  |
| **Mean July C.** | Temperature |  | X |  |  |  |  |
| **Mean August C.** | Temperature |  | X |  |  |  |  |
| **Mean September C.** | Temperature |  | X |  |  |  |  |
| **Mean October C.** | Temperature |  | X |  |  |  |  |
| **Mean November C.** | Temperature |  | X |  |  |  |  |
| **Mean December C.** | Temperature |  | X |  |  |  |  |
| **Prec. January C.** | Precipitation |  | X |  |  |  |  |
| **Prec. February C.** | Precipitation |  | X |  |  |  |  |
| **Prec. March C.** | Precipitation |  | X |  |  |  |  |
| **Prec. April C.** | Precipitation |  | X |  |  |  |  |
| **Prec. May C.** | Precipitation |  | X |  |  |  |  |
| **Prec. June C.** | Precipitation |  | X |  |  |  |  |
| **Prec. July C.** | Precipitation |  | X |  |  |  |  |
| **Prec. August C.** | Precipitation |  | X |  |  |  |  |
| **Prec. September C.** | Precipitation |  | X |  |  |  |  |
| **Prec. October C.** | Precipitation |  | X |  |  |  |  |
| **Prec. November C.** | Precipitation |  | X |  |  |  |  |
| **Prec. December C.** | Precipitation |  | X |  |  |  |  |
| **Annual mean** | Temperature |  |  | X |  |  |  |
| **Mean diurnal range** | Temperature |  |  | X |  |  |  |
| **Isothermality** | Temperature |  |  | X |  |  |  |
| **Temp. seasonality** | Temperature |  |  | X |  |  | 0.93 |
| **Max. Temp. of warmest month** | Temperature |  |  | X |  |  |  |
| **Min. Temp. of coolest month** | Temperature |  |  | X |  |  |  |
| **Temp. annual range** | Temperature |  |  | X |  |  |  |
| **Mean Temp. of wettest month** | Temperature |  |  | X |  |  |  |
| **Mean Temp. of driest month** | Temperature |  |  | X |  |  |  |
| **Mean Temp. of warmest quarter** | Temperature |  |  | X |  |  |  |
| **Mean Temp. of coldest quarter** | Temperature |  |  | X |  |  |  |
| **Annual mean of precipitation** | Precipitation |  |  | X |  | 0.62 |  |
| **Prec. of wettest month** | Precipitation |  |  | X |  |  |  |
| **Prec. of driest month** | Precipitation |  |  | X |  |  |  |
| **Prec. seasonality** | Precipitation |  |  | X |  |  |  |
| **Prec. of wettest quarter** | Precipitation |  |  | X |  | 0.49 |  |
| **Prec. of driest quarter** | Precipitation |  |  | X |  |  |  |
| **Prec. of warmest quarter** | Precipitation |  |  | X |  | 0.48 |  |
| **Prec. of coldest quarter** | Precipitation |  |  | X |  |  |  |

**Table 6**. LFMM median values per sampling site and environmental variables (LFFM input file: PCA scores and cold tolerance data (2).

| **Analysis** | **Sampling site** | **PC 1 = ENV1** | **PC 2= ENV2** | **PC 3 = ENV3** | **CT = ENV4** |
| --- | --- | --- | --- | --- | --- |
| PCA | CH200 | -533.73 | 230.69 | -42.07 | 6.40 |
|  | DH600 | -114.23 | -65.76 | 106.09 | 20.08 |
|  | DK800 | 46.29 | -386.85 | -50.737 | 23.00 |
|  | KT1300 | 601.67 | 221.92 | -13.28 | 11.86 |
| LFMM | - | 0.066 | 0.759 | 0.346 | -0.177 |

**Table 7**. Characteristic of amino acid before and after alternative base exchange at non-synonymous SNP position (3,4).

| **Chromosome** | **Position** | **Gene** | **Base** | **Alternative**  **base** | **Amino acid exchange** | **characteristic** |
| --- | --- | --- | --- | --- | --- | --- |
| NC_035107.1 | 59746123 | adenylate cyclase type 9* | G | T | P🡪H | nonpolar 🡪 polar (basic) |
| NC_035107.1 | 70557897 | proto-oncogene tyrosine-protein kinase ROS | T | C | I🡪V | nonpolar 🡪 nonpolar |
| NC_035108.1 | 223930033 | homeobox protein araucan | G | A | A🡪V | nonpolar 🡪 nonpolar |
| NC_035108.1 | 295810879 | uncharacterized protein LOC5566519* | G | A | E🡪K | polar (acidic) 🡪 polar (basic) |
| NC_035108.1 | 370218447 | breast cancer anti-estrogen resistance protein 3 | G | A | V🡪I | nonpolar 🡪 nonpolar |
| NC_035108.1 | 402025916 | toll-like receptor Tollo* | T | A | H🡪Q | polar (basic) 🡪 polar |
| NC_035109.1 | 278880294 | zinc finger CCCH domain-containing protein 13* | G | T | E🡪D | polar (acidic) 🡪 polar (acidic) |
| NC_035109.1 | 300326627 | uncharacterized protein LOC5574261 | T | A | I🡪K | nonpolar 🡪 polar(basic) |
| NC_035109.1 | 307981403 | probable peptide chain release factor C12orf65, mitochondrial* | A | T | F🡪I | nonpolar 🡪 nonpolar |
| NC_035109.1 | 308648742 | tubulin-specific chaperone D | G | A | V🡪I | nonpolar 🡪 nonpolar |
| NC_035109.1 | 314722675 | coatomer subunit beta' | G | A | A🡪T | nonpolar🡪 polar |
| NC_035109.1 | 319909869 | uncharacterized protein LOC5578603 | C | G | L🡪V | nonpolar🡪 nonpolar |

**Table 8.** Significantly enriched GO terms (p < 0.05) among candidate genes (EAP-OW) per ENV and their biological functions involved in climate adaptation. To increase resolution, the GO term analysis was conducted per ENV (ENV1 ~ altitude, ENV2 ~ precipitation, ENV4 = cold tolerance) as described by (5). The definition of GO terms was used from the webpage: (6). If literature for comparison was accessible, the biological function and the link with ENVs was discussed.

| **Enriched GO term** | **ENV1** | **ENV2** | **ENV4** | **Definition of GO terms involved in climate adaptation** | **Biological function involved in climate adaptation** |
| --- | --- | --- | --- | --- | --- |
| small GTPase mediated signal transduction (GO:0007264) | X | X | X | Any series of molecular signals in which a small monomeric GTPase relays one or more of the signals. |  |
| protein phosphorylation (GO:0006468) | X | X | X | The process of introducing a phosphate group on to a protein. | One of the most important post-translational modifications is protein phosphorylation. Protein phosphorylation is vital for the coordination of organic and cellular function such as the regulation of metabolism, proliferation, apoptosis, subcellular trafficking, inflammation, and other important physiological processes (7). ENV1, ENV2 and ENV4 seem to affect this important function. This GO term correlating with climate variation was also enriched in a GEA study in ants (8), whereas in the GEA study of Waldvogel (5) this GO term was enriched in candidate genes for local adaptation. |
| transmembrane receptor protein tyrosine kinase signaling pathway (GO:0007169) | X | X | X | A series of molecular signals initiated by the binding of an extracellular ligand to a receptor on the surface of the target cell where the receptor possesses tyrosine kinase activity, and ending with regulation of a downstream cellular process, e.g. transcription. | The major mechanism for intercellular communication during development as well as in the adult organism and in disease-associated processes is the signaling through receptor tyrosine kinases (9). ENV1, ENV2 and ENV4 seem to affect this important function. This GO term correlating with climate variation was also enriched in a GEA study in ants (8). |
| ubiquitin-dependent protein catabolic process (GO:0006511) | X |  | X | Chemical reactions and pathways leading to a breakdown of a protein or peptide by hydrolysis of its peptide bonds. |  |
| translational termination (GO:0006415) | X |  | X | The process that leads to the release of a polypeptide chain form the ribosome when a termination codon on the mRNA occurs. |  |
| regulation of pH (GO:0006885) |  | X |  | Within an organism or cell, processes involved in the maintenance of an internal equilibrium of hydrogen ions, thereby modulating the internal pH. | This GO term is upregulated in the highly secreting pleuropodia. Pleuropodia appear on the first abdominal segment in embryos of insects. This organ secrete a “hatching enzyme” that enables the hatching of larvae by digesting the serosal cuticle (10). Thus, the association of this GO term with precipitation adds up. |
| sodium ion transport (GO:0006814) |  | X |  | Movement of sodium ions within, into or out of a cell, or between cells by some agents such as a transporter or pore. | In *Ae. aegypti* larvae the exchange of sodium occurs through the anal papillae, as well as smaller amounts enter the haemolymph through the gut and general body surface (11). In Nepal, sodium as well as chloride concentration in rainfall is high, especially in the monsoon season (12). Thus, larvae may need to cope with concentrations of these ions, which may affect survival and explain the association with precipitation. |
| Proteolysis (GO:0006508) |  | X | X | Hydrolysis of proteins into smaller polypeptides and/or amino acids, respectively by cleavage of their peptide bonds. | Temperature extremes affect protein stability and cells have to cope with stress-induced protein denaturation (13). Adaptation of the proteolysis machinery implies a response to the upper temperature extremes along the thermal cline (5). In the GEA study of Waldvogel (5)within *Chironomidae* species, this GO term was enriched in candidate genes for climate adaptation and was associated with warm temperatures, whereas in this study the GO term is associated with precipitation and cold tolerance. In addition, this GO term was significantly enriched among candidate genes of climate differentiation in *Drosophila* (5,14). |

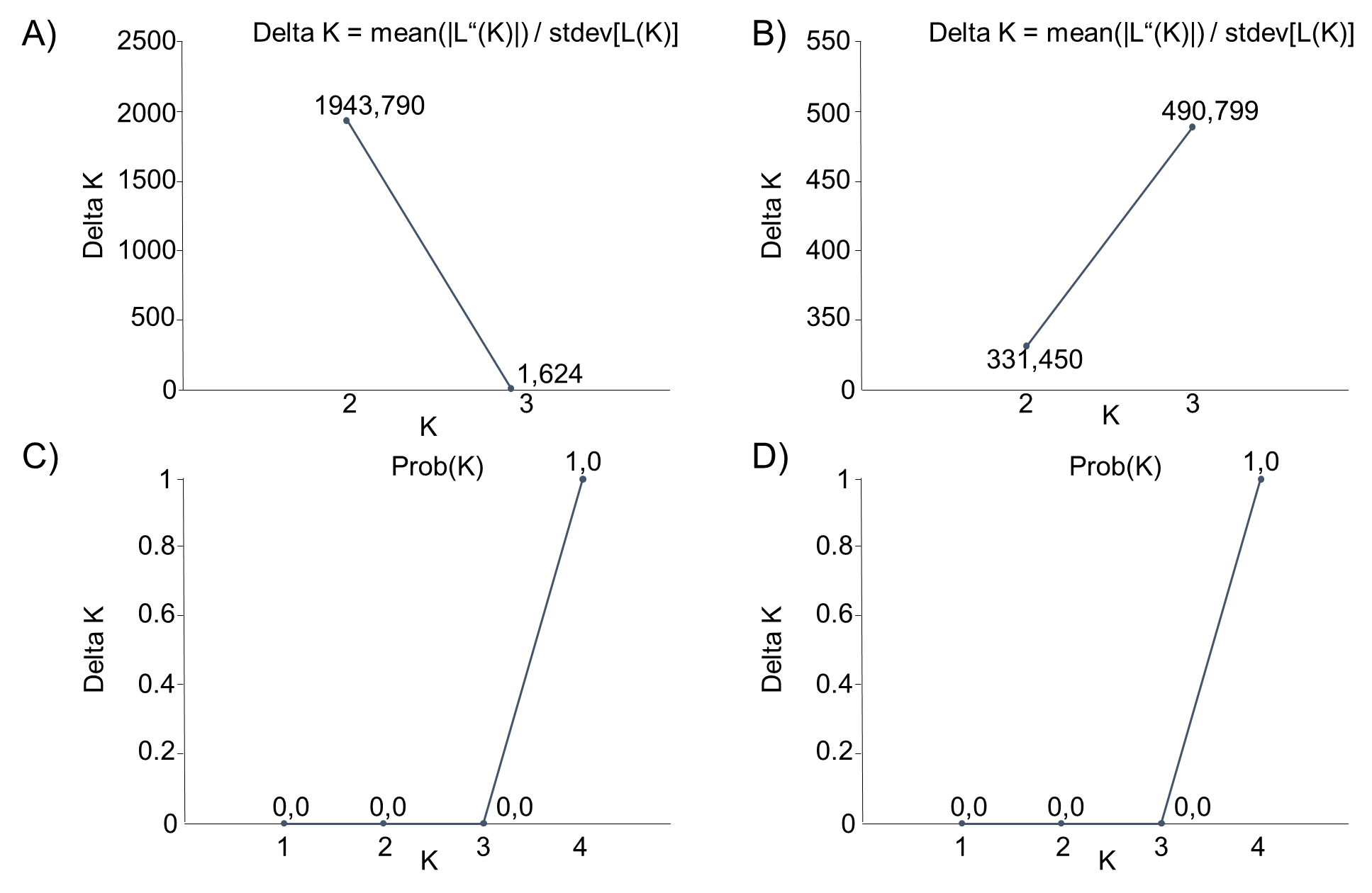

**Figure 1**. Delta K (A, B) and Probability by K (C,D) from the structure analysis. Delta K=Optimal K by Evanno is given. A+C) show the result of the STRUCTURE/CLUMPAK comparison between Nepal and Africa, Costa Rica and Australia using 6 microsatellite regions (15). Division of runs by mode for K=2 was 10/10 and for K=3 9/10, 1/10. C+D) show the results of the STRUCUTRE/ CLUMPAK comparison of only Nepalese populations using 11 microsatellite regions. Division of runs by mode for all K1-4 was 10/10.

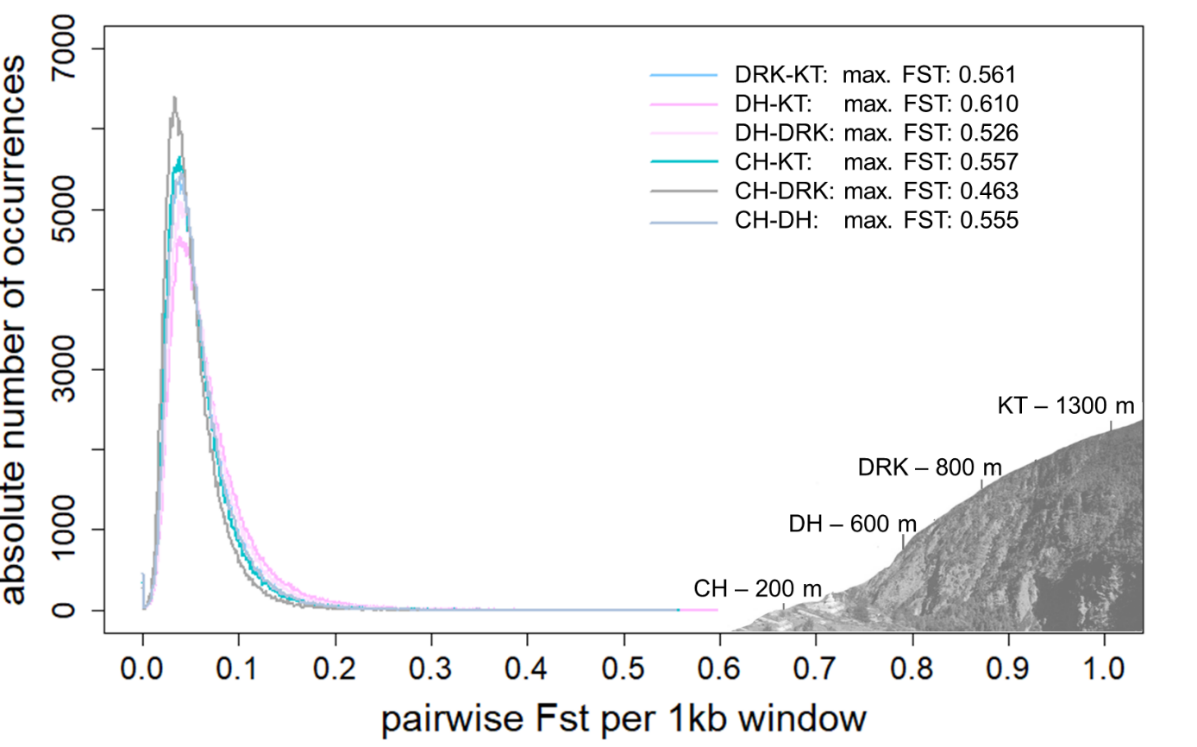

**Figure 2**. Pairwise F_ST_ distribution per 1 kb-windows (OW) of Nepalese *Ae. aegypti* populations sampled along an altitudinal gradient in Nepal (200-1300m). Altitude of sampling sites of *Ae. aegypti* populations in Central Nepal: CH200 = 200 m asl (Chitwan), DH600 = 600 m asl (Dhading), DK800 = 800 m asl (Dharke), KT1300 = 1300 m asl (Kathmandu).

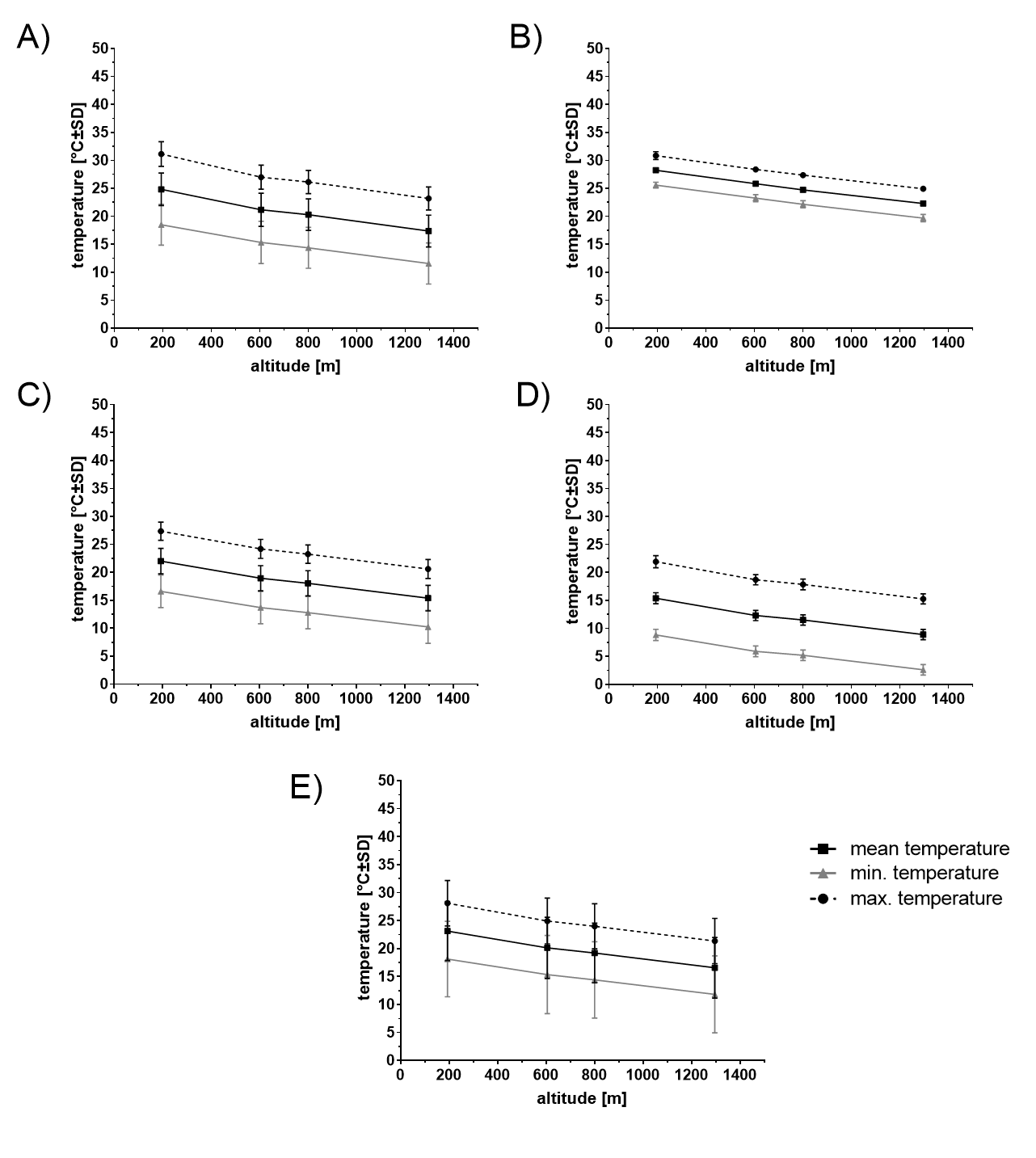

**Figure 3.** Climate (air temperature) along the altitudinal gradient in Central Nepal. Mean, minimum and maximum temperature (CHELSA data) in different time seasons from 1979 to 2013: A) pre-monsoon (March-May), B) monsoon (June-September), C) post-monsoon (October-November), D) winter (December-February), E) annual.

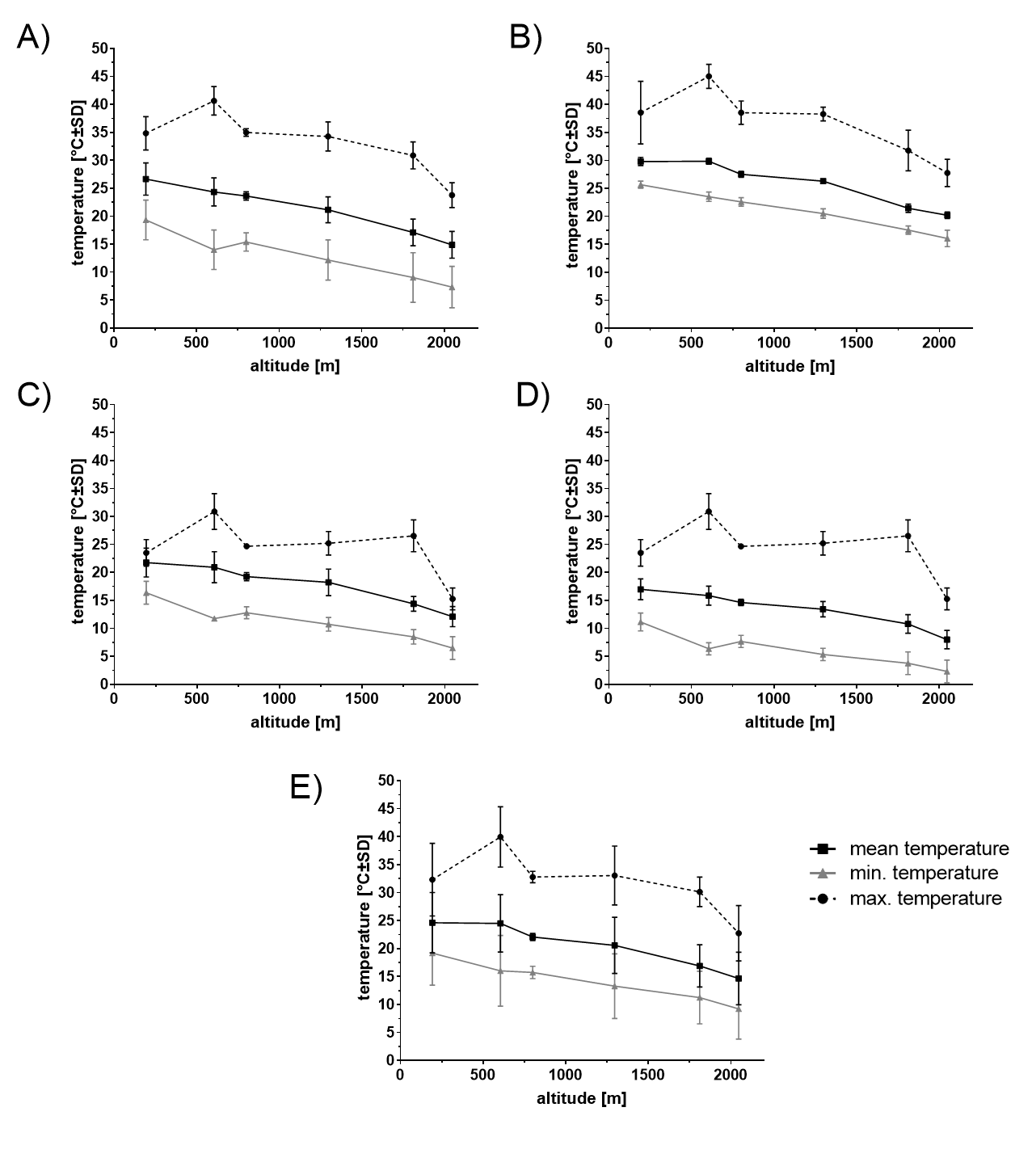

**Figure 4**. Microclimate (air temperature) along the altitudinal gradient in Central Nepal. Mean, minimum and maximum temperature (HOBO logger data; microclimate see also Additional File 1 Table 3) in different seasons from November 2017 to March 2019: A) pre-monsoon (March-May), B) monsoon (June-September), C) post-monsoon (October-November), D) winter (December-February), E) annual. The 800 meter population was interpolated using raw data of loggers.

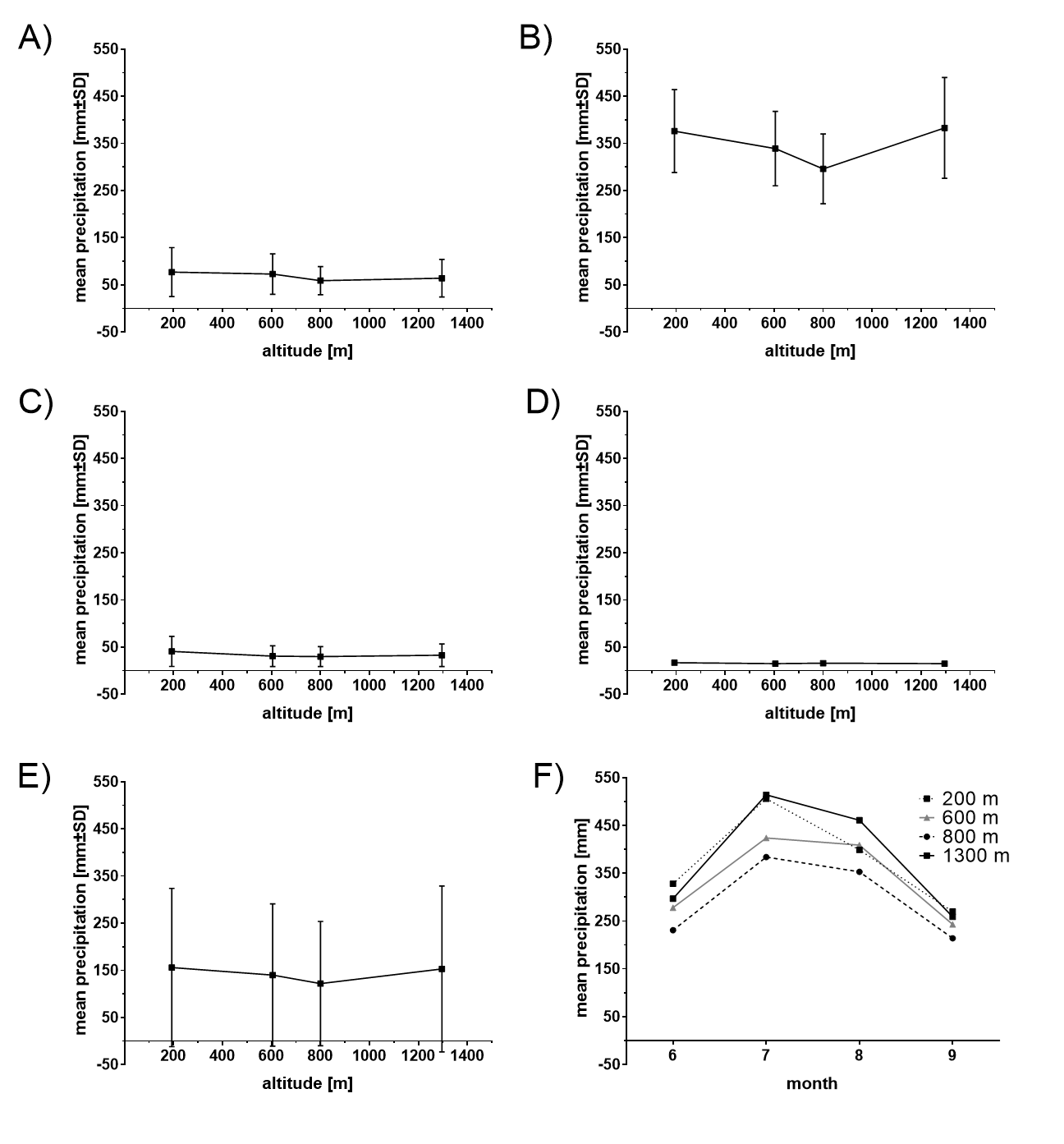

**Figure 5.** Precipitation along the altitudinal gradient in Central Nepal. Mean precipitation (CHELSA data) in different seasons from 1979 to 2013: A) pre-monsoon (March-May), B) monsoon (June-September), C) post-monsoon (October-November), D) winter (December-February), E) annual, F) monthly monsoon mean precipitation

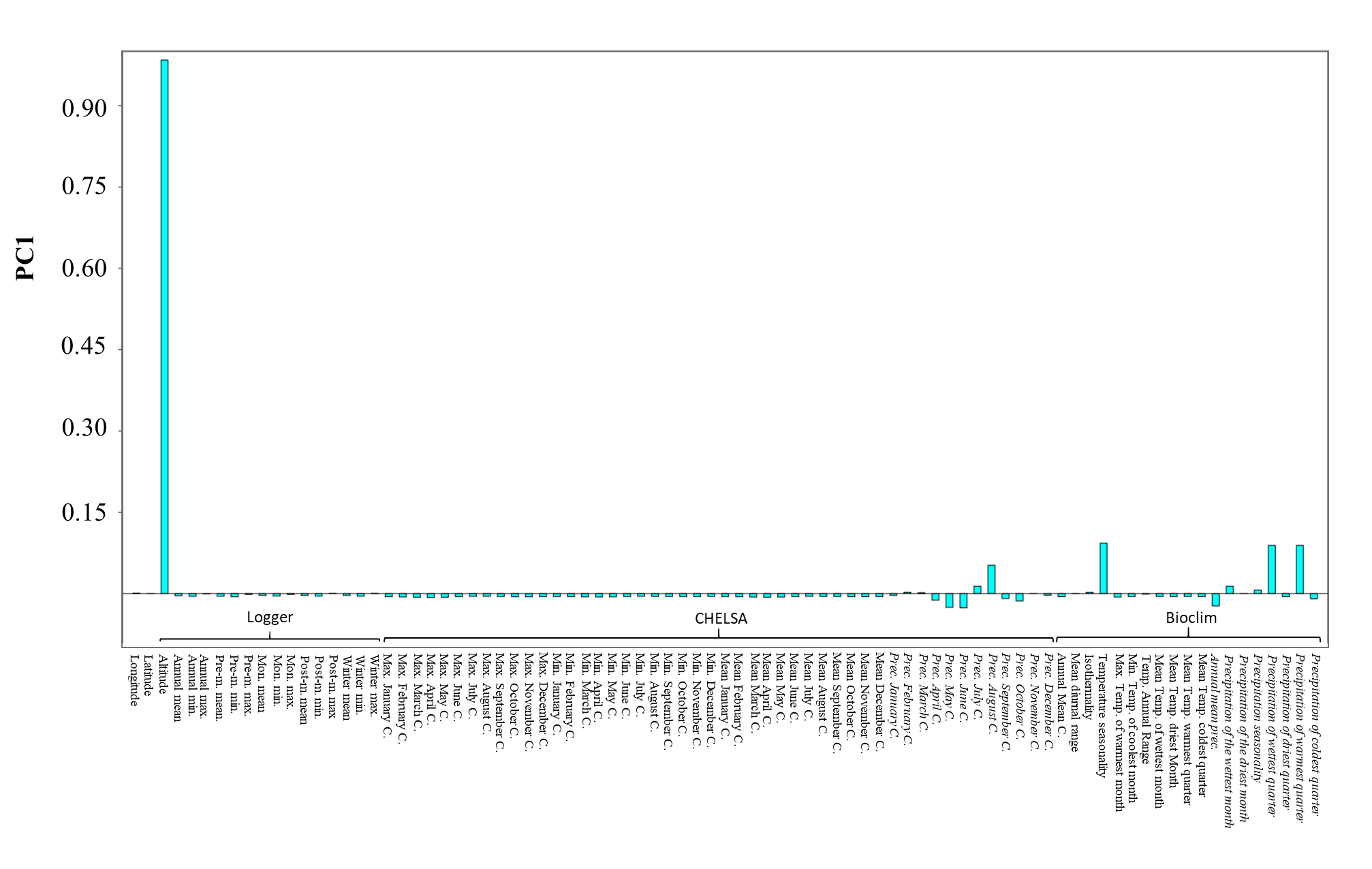

**Figure 6.** Loadings from PC (principal component) analysis: PC1 is associated with altitude. Precipitation related variables are highlighted using Italic font. Temperature related variables and longitude, latitude and altitude are given in non-italic font.

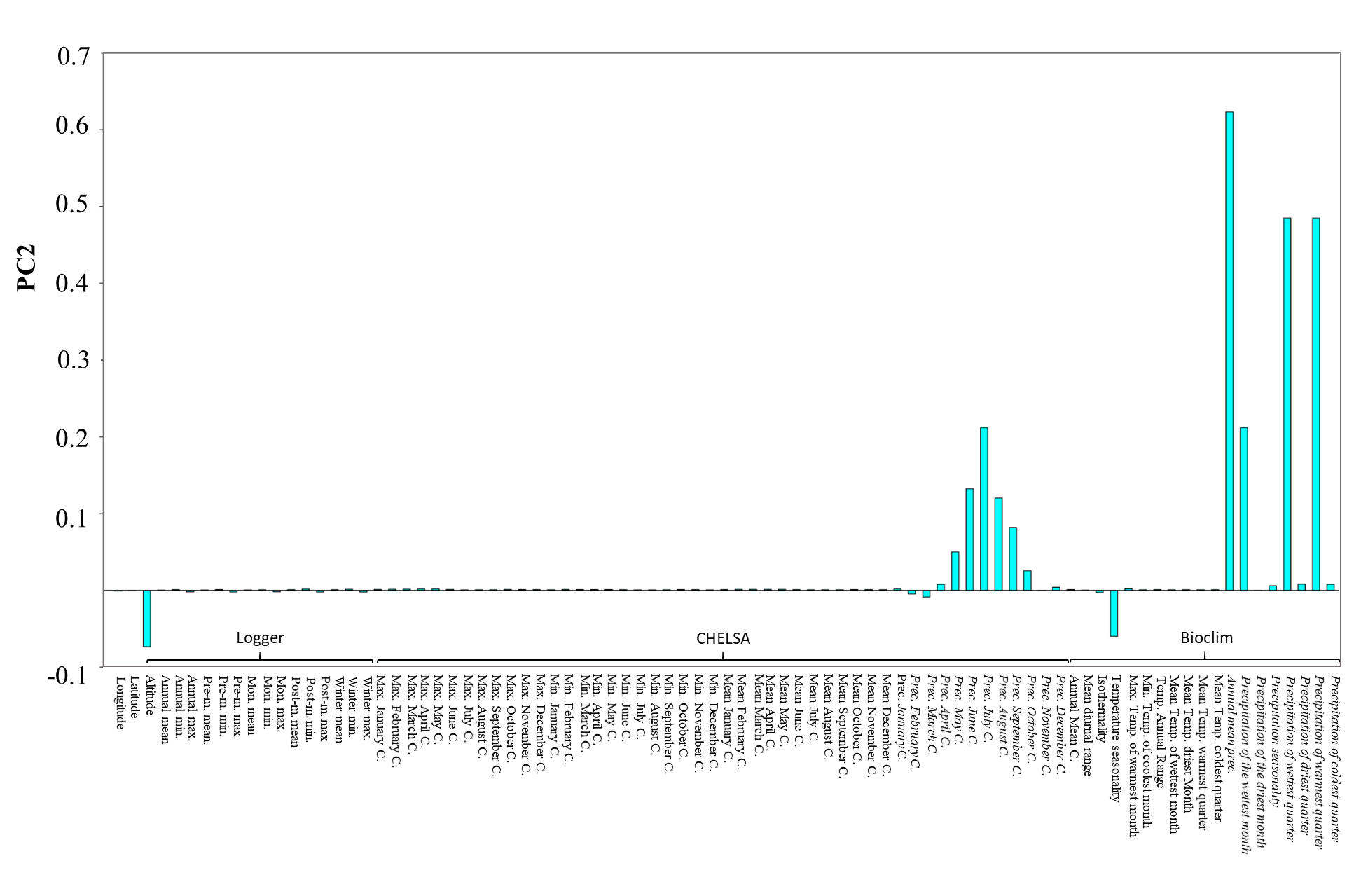

**Figure 7.** Loadings from PC (principal component) analysis: PC2 is associated with precipitation. Precipitation related variables are highlighted using Italic font. Temperature related variables and longitude, latitude and altitude are given in non-italic font.

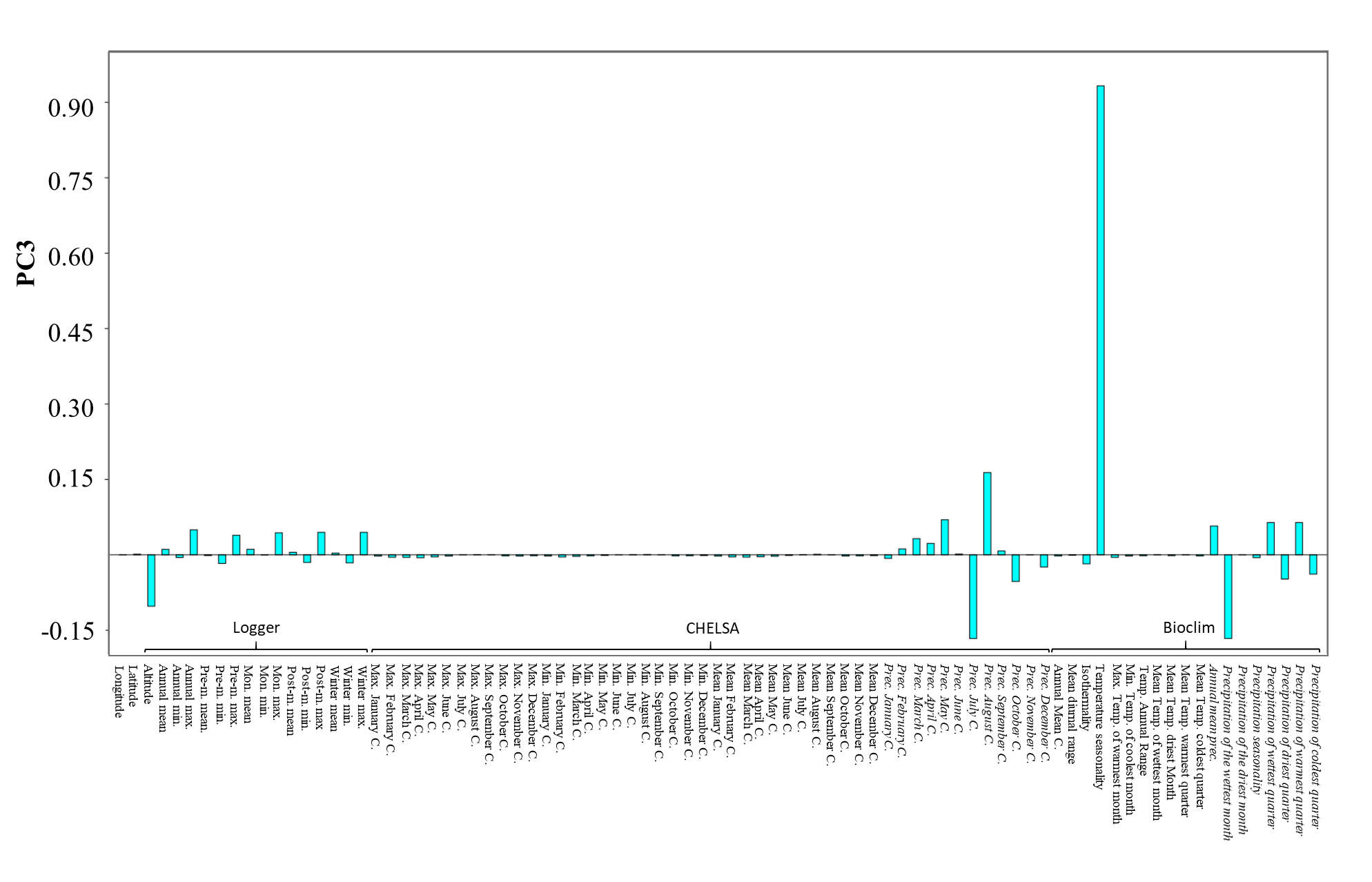

**Figure 8.** Loadings from PC (principal component) analysis: PC3 is associated with seasonality. Precipitation related variables are highlighted using Italic font. Temperature related variables and longitude, latitude and altitude are given in non-italic font.

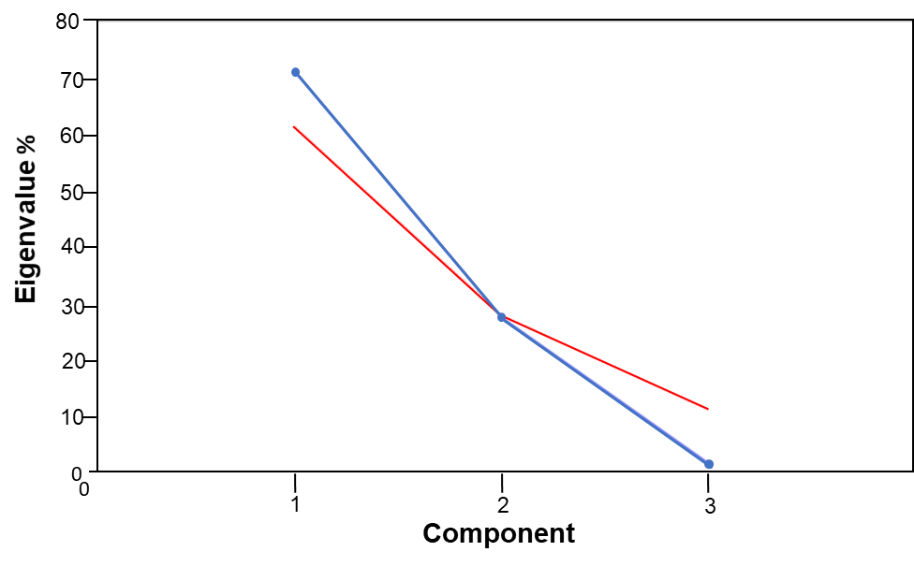

**Figure 9.** Distribution of Eigenvalues (%) of principal components (blue line). Gained from principal component analysis with 85 climatic variables (logger data, CHELSA, Bioclim) at 4 different sampling sites of *Ae. aegypti* populations along an altitudinal gradient in Nepal. Broken stick analysis is given as red line and indicates that PC3 componenent under this line is non significant.

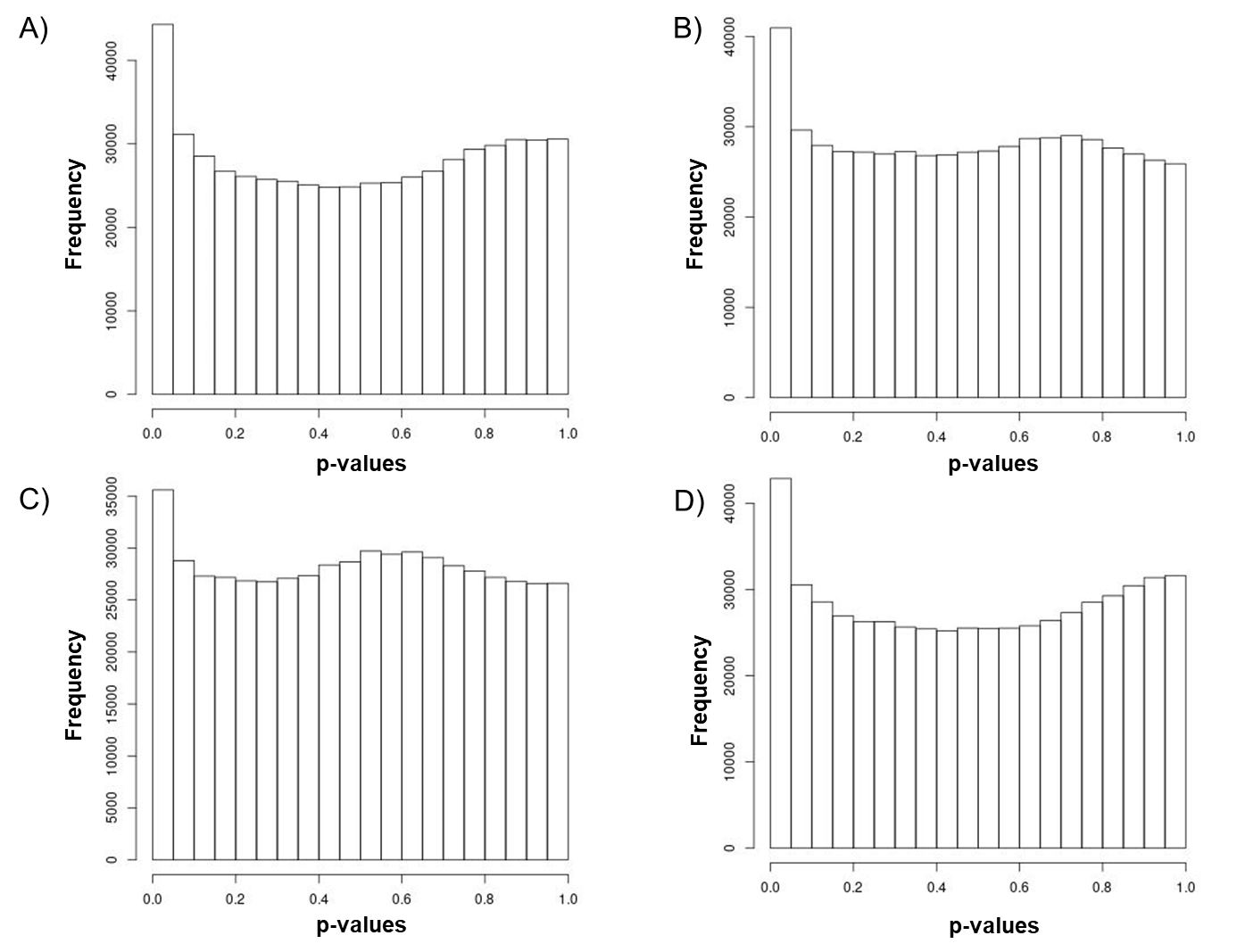

**Figure 10.** The frequency distribution of adjusted p-values after association (genotype-enviornment association analysis) to four different environmental variables (ENVs) using LFMM. 1) ENV1 = PC1 associated with altitude, 2) ENV2 = PC2 associated with precipitation, 3) ENV3 = PC3 associated with seasonality and 4) ENV4 = cold tolerance of *Ae. aegypti* (2).

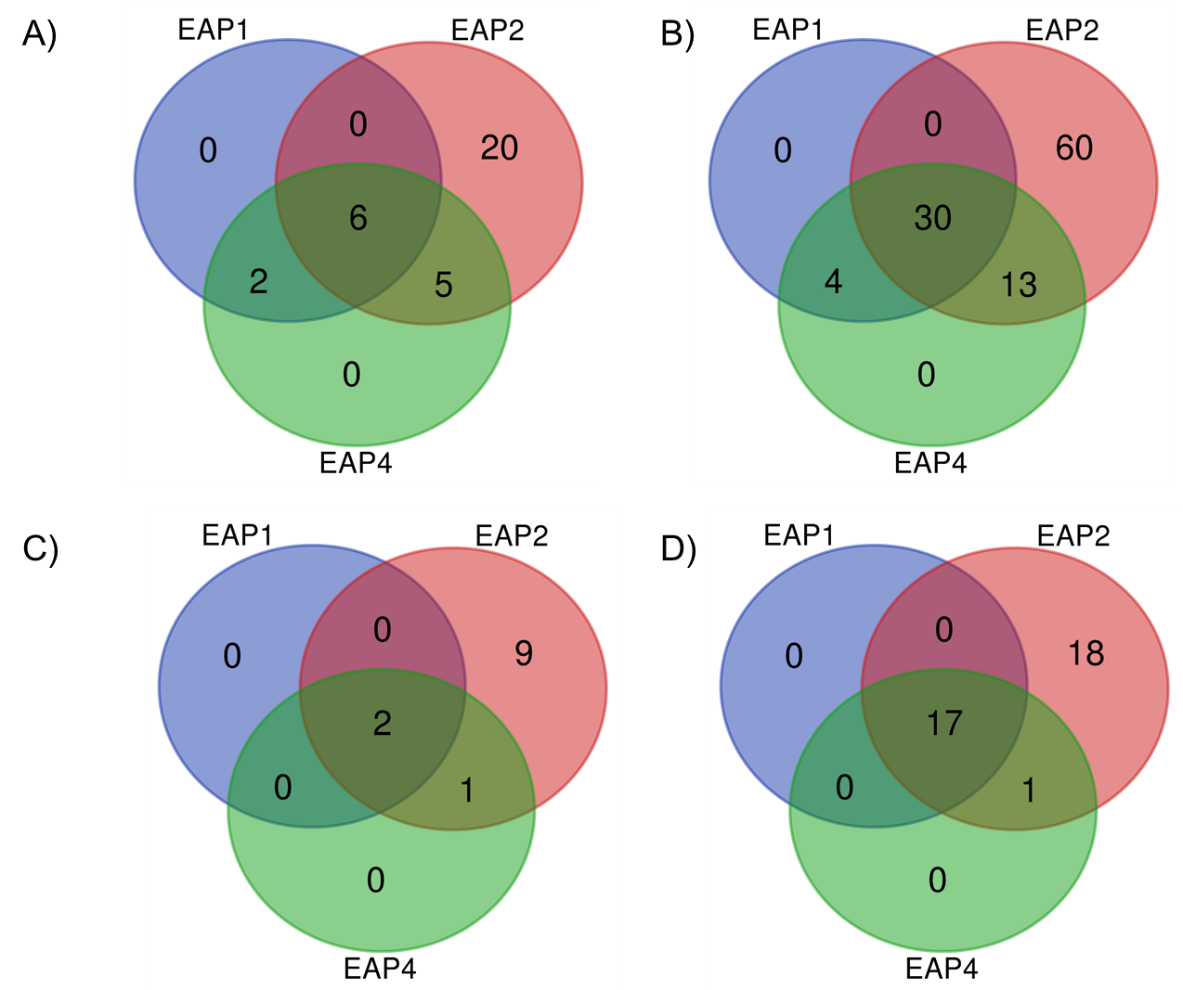

**Figure 11.** Gene IDs (A, C) or protein IDs (B,D) present in all different significant environmental variable associated positions (EAP) laying in an overlapping singificant 1kb-F_ST_-window (OW, A and B) and contain a non-synonymous mutation (C and D).

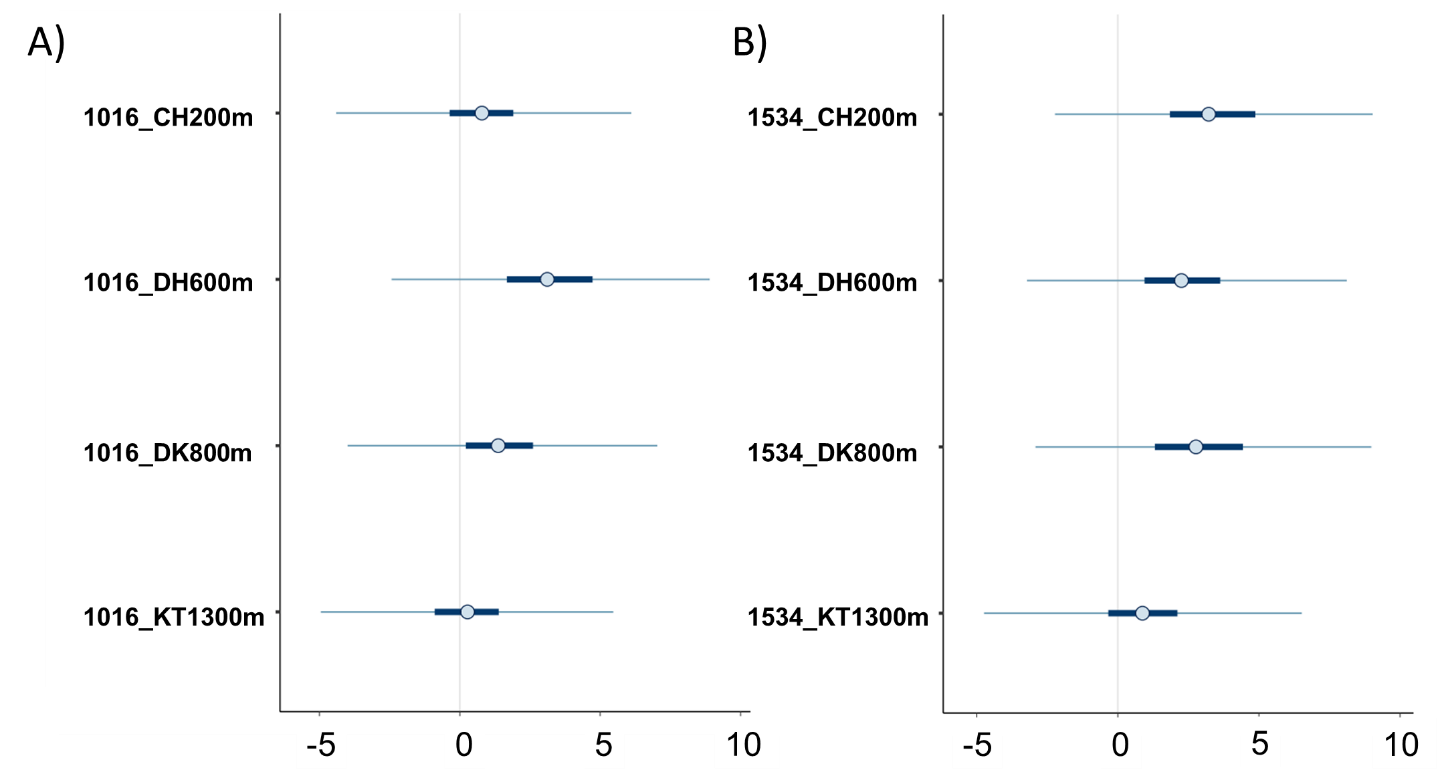

**Figure 12.** Posterior uncertainty intervals for kdr mutation a) V1016G and b) F1534C. Depicted are the median, 50% and 90% posterior intervals. CH200 = 200 m asl, DH600 = 600 m asl, DK800 = 800 m asl, KT1300 = 1300 m asl.

**Supplementary Information 1**

**Go terms**

In the present study, the GO term ‘transmembrane receptor protein tyrosine kinase signaling pathway‘ and ‘protein phosphorylation’ were found to be associated with all environmental clines. The first plays a major role in the intercellular communication during the development (9). The second is important for the regulation of organic and cellular functions such as the metabolism, proliferation or apoptosis (7), underlining the impact of the environmental clines on the life-cycle of *Ae. aegypti*. Further, both of these GO terms were already found to be enriched in candidate genes for climate variation in ants (8). In addition, ‘proteolysis’ is associated with precipitation and cold tolerance and may imply a response to climate extremes/stress (5,14). The GO term ‘proteolysis’ was already found to be enriched in candidate genes for climate adaptation in other GEA studies in *Chironomus* and *Drosophila* (5,14). The GO term ‘regulation of pH’, which is associated with precipitation, was already previously identified for playing a role in the hatching of larvae (10). The association with precipitation aligns with the fact that rainwater is needed for the hatching of *Ae. aegypti* eggs (16). In Nepal, sodium concentration in the rain water is high, and especially in the monsoon season (12). Thus, immature life stages may need to cope with varying concentrations of these ions which explains the association with precipitation of the GO term ‘sodium ion transport’.

**Candidate genes containing non-synonymous mutations**

The candidate gene ‘proto-oncogene tyrosine-protein kinase ROS’ including a non-synonymous mutation is associated with ENV1, ENV2 and ENV4 (altitude, precipitation and CT survivorship) is suggested to play a role in development or/and energy metabolism (17,18). These biological processes are important in survival after cold temperatures (CT survivorship), the start of a life-cycle as well as the general life-cycle linked to precipitation and is influenced by the environment at different altitudes.

Precipitation is a vital factor for *Ae. aegypti* (16). In Nepal vector and dengue occurrence is linked with precipitation or more specifically the monsoon season (19–21). Thus, the following genes which play a role in immune response were all associated with precipitation. The ‘toll-like receptor Tollo’ was already studied in *Ae. aegypti* and plays a role in the immune response especially in the anti-dengue defense (22–28). In addition, the non-synonymous mutations within this candidate gene leads to an amino acid with different characteristics (Additional File 1 Table 7). Other candidate genes play also a role in dengue viral defense (immune response) such as the ‘probable peptide chain release factor C12orf65’ (also plays a role in protein regulation- Table 4), ‘breast cancer anti estrogen resistance protein 3’ (29) as well as the ‘zinc finger CCCH domain-containing protein 13’ (30,31). With increasing altitude, the dengue risk decreases with being the greatest below 500 m asl and moderate between 550-1500 m asl (32). This altitudinal distribution of dengue disease risk could explain the association between altitude and the ‘breast cancer anti estrogen resistance protein 3’ that is as well involved in dengue infection or defense (29).

The candidate gene ‘adenylate cyclase type 9’ is associated with precipitation (ENV2) was the only gene in the climate adaptation analysis that is in parallel involved in insecticide resistance and contains a non-synonymous mutation leading to a characteristic amino acid exchange. In the mosquito *Culex quinquefasciatus,* this gene is involved in the regulation of the resistance-related P450 gene expression (33,34). In Nepal, insecticides use is declining as summarized by (35)The association of an insecticide resistance related gene to precipitation might be justified by the use of insecticides mainly in the monsoon season (high peak mosquito season) and the influence of rainfall on the distribution of insecticides (36), subsequently this influences the vector *Ae. aegypti*.

Candidate genes involved in the life-cycle such as the ‘Tubulin-specific chaperone D’, ‘coatomer subunit beta’, ‘homeobox protein auracan’ and the ‘zinc finger CCCH domain-containing protein 13’ are likewise associated with precipitation (ENV2; Table 4). ‘Tubulin-specific chaperone D’ is involved in the reproduction (37). The ‘coatomer subunit beta’ including a non-synonymous mutation leading to a characteristic amino acid shift has an major impact on blood digestion and egg maturation (38,39). The ‘homeobox protein auracan’ is involved in the embryonic development (40,41) and the ‘zinc finger CCCH domain-containing protein 13’ has an impact on development processes and reproduction (30,31,42). The association with precipitation of these candidate genes underlines the importance of precipitation in the life cycle of *Ae. aegypti*.
